# Using large language models to study human memory for meaningful narratives

**DOI:** 10.1101/2023.11.03.565484

**Authors:** Antonios Georgiou, Tankut Can, Mikhail Katkov, Misha Tsodyks

## Abstract

One of the most impressive achievements of the AI revolution is the development of large language models that can generate meaningful text and respond to instructions in plain English with no additional training necessary. Here we show that language models can be used as a scientific instrument for studying human memory for meaningful material. We developed a pipeline for designing large scale memory experiments and analyzing the obtained results. We performed online memory experiments with a large number of participants and collected recognition and recall data for narratives of different lengths. We found that both recall and recognition performance scale linearly with narrative length. Furthermore, in order to investigate the role of narrative comprehension in memory, we repeated these experiments using scrambled versions of the presented stories. We found that even though recall performance declined significantly, recognition remained largely unaffected. Interestingly, recalls in this condition seem to follow the original narrative order rather than the scrambled presentation, pointing to a contextual reconstruction of the story in memory.

## 1 Introduction

In the classical paradigm for studying human memory, participants are presented with randomly assembled lists of words and then perform memory tasks such as recognition and recall (see review in [Kahana, 2020]). A wealth of results has been obtained in these studies. For instance, it has been found that words at the end and the beginning of the list have a higher chance of being recalled (recency and primacy effects, respectively), and there is a tendency to recall words close to each other in the list (contiguity, [Kahana, 1996]). Moreover, it was found that when the presented lists grow in length, even though the average number of recalled words (*R*) is increasing, a progressively smaller fraction of the words is recalled [Murdock Jr, 1962]. Several authors have addressed the issue of the mathematical form of the dependence of *R* on list length and found that the best description for this dependence is provided by power-law relations, *R* ∼*L*^*α*^, with exponents *α* generally below one [Murray et al., 1976]. It is well known that recall also depends on multiple experimental factors such as e.g. the presentation rate of words, the age of the participants, etc. However, in recent work, some of the authors discovered that if recall performance is analyzed as a function of a number of *remembered* (*M*), rather than presented words, the relation becomes universal and is described by the analytical form: 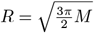 [Naim et al., 2020]. Moreover, this relation follows from a simple deterministic model where words are retrieved one by one according to a random symmetric matrix of ‘similarities’ reflecting their long-term encoding in memory, until the process enters a cycle and no more words can be recalled. The number of remembered words *M*, itself can be predicted by the *retrograde interference* model that assumes that each new word erases some of the previously presented words according to the ‘valence’ or ‘importance’ of each word [Georgiou et al., 2021, 2023].

While it is remarkable that human memory for random material can be described with universal mathematical relations, it is of course much more important and exciting to try to understand how people remember more natural, meaningful information. After the pioneering work of Bartlett [1932], many studies considered recalls of narratives. As opposed to random lists, narratives convey meaning, and hence have structure on multiple levels which influences recall, as was confirmed in many previous publications (see Section 3). The first challenge in understanding narrative recall is the fact that people tend not to recall the narrative verbatim. Rather, they remember what the narrative is about and retell it in their own words [Gomulicki, 1956; Fillenbaum, 1966; Sachs, 1967]. Counting correctly recalled words is therefore not a good score of recall, and the better score used in many studies, that we also adopt in our work, is a count of recalled ‘ideas’, or ‘clauses’ (see e.g. [Bransford and Johnson, 1972]). Using this method however requires a human-level of understanding of narratives and recalls, making collecting large amounts of data difficult and extremely time-consuming to analyze. In our study, we develop a way to overcome this and other difficulties by using large language models (LLMs) to assist in the analysis and design of experiments, as described later. In particular, we use LLMs to generate new narratives of a particular type and length, and to score human recalls obtained in multiple experiments performed over the internet. In addition to recall, we also performed *recognition* experiments (where people are requested to indicate whether a specific clause was in the presented story or not) in order to estimate how many clauses people remember after reading the narrative. To this end, we use LLMs to generate plausible lures, i.e. novel clauses that could have potentially appeared in the narrative.

Inspired by our previous results, we wanted to understand how recognition and recall performance scale up with narrative length as it increases and what the relation between them is. To this end, we performed a large number of experiments over the internet using the Prolific platform (www.prolific.com). We also compared the recall and recognition performance of original narratives with their scrambled versions in order to elucidate the effects of comprehension on different aspects of memory. Since there are different types of narratives that could potentially be more or less difficult for people to remember and recall, we decided to focus on one particular type of narrative first studied in the famous paper by Labov and Waletzky [1966] that established the field of narratology, namely the oral retelling of personal experience, told by real people, and the variants of those generated by LLMs (see later). While being collected in a research setting, these spoken recollections of dramatic personal episodes are close to the natural way people share their experiences in real life and therefore are of special interest for studying human memory.

## 2 Results

### 2.1 LLM-assisted recall and recognition experiments

For the purpose of this study, we have chosen several narratives of different lengths from Labov [2013] and Labov and Waletzky [1966]. As part of the analysis in these publications, narratives were segmented into an ordered set of *clauses*, which are “the smallest unit of linguistic expression which defines the functions of narrative” [Labov and Waletzky, 1966]. In other words, they are the smallest meaningful pieces which still serve some function in communicating a narrative. Since these are spontaneous narratives spoken in local dialect, they are characterized by a number of features which are awkward to transcribe (pauses, repetition, gestures) as well as non-standard (and sometimes outdated) English vernacular. These factors complicate comprehension when participants have to read narratives on the computer screen. We therefore instructed LLMs to generate new narratives modeled on the original ones, i.e. exhibiting a similar type of a event sequence and the overall length in terms of the total number of clauses. In particular, the LLM-generated narratives inherited the segmentation from the original story, i.e. the number of clauses was the same and the information contained in the corresponding clauses had a similar role in their respective narrative (see Methods Section 5 for details of narrative generation, and some examples in Appendix A.1). Eight narratives were selected for subsequent memory experiments, ranging from 18 to 130 clauses in length. We presented each narrative to a large number of participants (∼ 200) who then performed either recall or recognition tasks. In the subsequent analysis, we treated clauses as the basic units that together communicate the meaningful information contained in the narrative. In particular, we quantified each individual recall by identifying which of the clauses in the narrative were recalled, determining this by whether information contained in this clause is present in the recall. We simplify the analysis by considering each clause as being either recalled or not. This scoring of recalls is traditionally performed by human evaluators and is very time-consuming. We, therefore, *prompted* an LLM to define which of the clauses of the original narrative were recalled and in which order. Here we utilized the remarkable ability of modern LLMs to respond to instructions, provided as *prompts* written in standard English (as opposed to a programming language), to perform novel tasks without any additional training (known as zero-shot prompting or ‘in-context’ learning [Brown et al., 2020], see Appendix A for more details).

To test the ability of the LLM to adequately score human recall (with appropriate prompting as described in the Methods Section 5 and Appendix A), we performed an additional set of recall experiments with a specially LLM-generated narrative and compared the LLM-performed recall scoring to the one conducted manually by the authors (see Methods Section 5 for details). To this end, we calculated the fraction of participants who recalled each particular clause, i.e. the clause’s recall probability (*P*_*rec*_), as judged by the LLM and by the authors. For nearly all of our analysis, the LLM we used was OpenAI’s GPT-4 (see Appendix B.2 for comparison between different LLMs). As shown in Figure 1, GPT-4 scoring of recalls results in recall probabilities close to ones obtained by human evaluations for a great majority of the clauses. Moreover, variability of scoring is comparable between GPT-4 and human evaluators (compare Figure 1B and C). Interestingly, the LLM has a greater correlation with the mean human scoring (r = 0.94) than with any individual scoring (r = 0.92, 0.90, 0.90) (see Table 1 in Appendix B; c.f. Michelmann et al. [2023]).

**Table 1:**
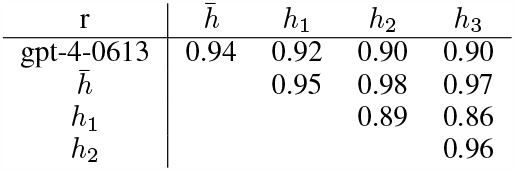
A comparison between gpt-4, mean human 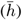, and individual human scorers (*h*_1_, *h*_2_, *h*_3_). Each entry in the table gives the correlation coefficient (r-value) between the corresponding row and column label.

**Table 2:**
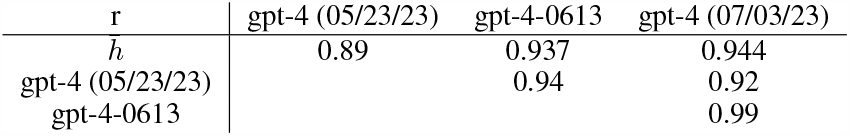
A comparison on gpt-4 scoring compiled at different times of the year. Evidently, the model improves over time in approximating the mean human scoring, with the most recent model achieving a correlation closest to one. The entries of the table show correlation coefficients (r-values) between the corresponding row and column variables.

| r | gpt-4 (05/23/23) | gpt-4-0613 | gpt-4 (07/03/23) |
|---|---|---|---|
| *h̄* | 0.89 | 0.937 | 0.944 |
| gpt-4 (05/23/23) |  | 0.94 | 0.92 |
| gpt-4-0613 |  |  | 0.99 |

**Figure 1.**
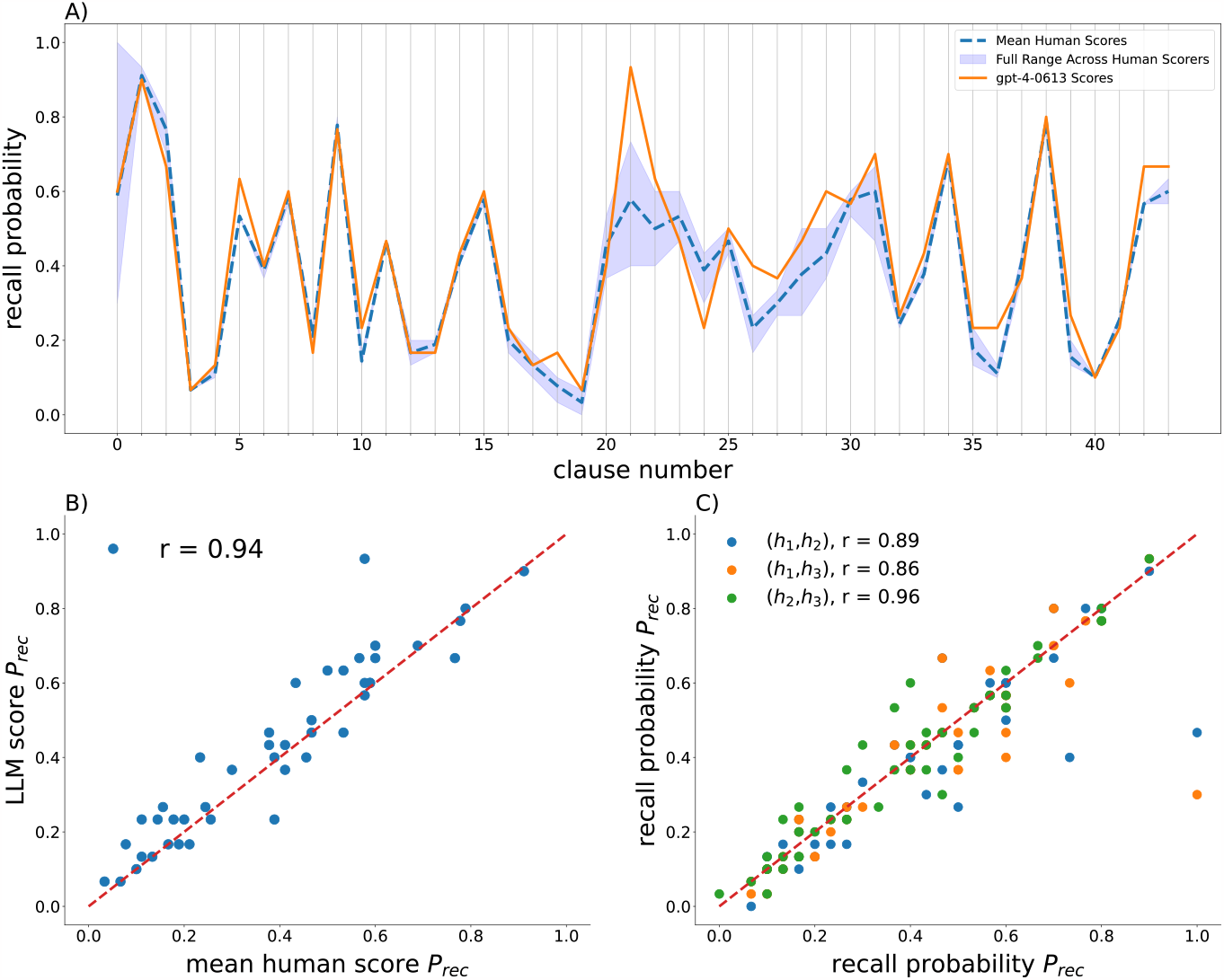
Reliability of LLM scoring of recalls. 3 authors and GPT-4 (OpenAI API model `gpt-4-0613`) performed scoring of 30 recalls by answering the question of whether the information present in each particular clause is present in each individual recall. (A): comparison between recall probabilities *P*_*rec*_ for each clause, as calculated from GPT-4 scores (orange) and average human scores (dashed blue). The full range of human scores is given by the shaded blue region. (B): A strong correlation between human and GPT-4 scores across clauses, with a correlation coefficient (r-value) of 0.94. (C): Correlations between individual human scores, shows overall a strong agreement between human scorers.

One prominent feature seen in Figure 1A is a wide range of recall probabilities for different clauses. In other words, while some clauses are recalled by most of the participants, other clauses are not. Such a wide distribution of *P*_*rec*_’s across the clauses was observed in recalls of all the narratives and contrasts sharply with corresponding results for random lists of words where *P*_*rec*_’s are rather uniform except for the ones in the beginning and the end of the list (see Appendix D). This wide distribution of *P*_*rec*_s is apparently due to the fact that not all clauses have similar importance for communicating the narrative. Indeed if we select the clauses with high enough *P*_*rec*_, we usually get a good summary of the narrative (see Appendix E for examples).

We also performed recognition experiments in order to estimate the average number of clauses that participants remember after presentation. As we explain below, this analysis requires a large number of plausible *lures*, i.e. novel clauses that could have possibly been in the narrative and hence cannot be easily distinguished from the true clauses using context and style. Generating these lures is highly nontrivial as it requires an understanding of the narrative. This makes manually generating lures very challenging and time-consuming, which is why we utilized LLMs for this purpose (see Appendix A.2 for prompts and example output). Using the LLM, for each story we obtained the same number of lures as true clauses, and sampled 10 clauses from this entire pool of 2L clauses uniformly and randomly for the testing phase. Participants would then see one clause at a time, and were asked whether the clause was in the presented narrative or not. We checked that presenting several clauses for recognition does not result in systematic drift in performance, i.e. no output interference was detected in our experiments (Criss et al. [2011]; see Appendix C). We then estimated the number of clauses retained in memory after presentation of the narrative (*M*) from the fraction of ‘hits’, i.e. correct recognitions of the true clauses (*P*_*h*_) and the fraction of ‘false alarms’, i.e. reporting lures as true clauses (*P*_*f*_). In particular, we assume that if the participant remembers a given clause, they always recognize it as being part of the narrative; otherwise they still give a positive answer with probability *P*_*f*_. The total probability of a correct recognition is then given by

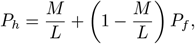

from which we obtain

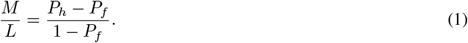

This equation emphasizes the importance of using lures in recognition experiments, since without lures we would have an uncontrolled tendency of participants to indicate any presented clause as a true one irrespective of whether or not they remember it.

### 2.2 Scaling of Recall and Memory

Having each narrative seen by roughly 200 participants, half of them doing recall and another half recognition, we were able to determine the average number of remembered clauses (*M*) and recalled clauses (*R*). As expected both *M* and *R* grow with the length of the narrative presented, as measured by the number of clauses in the narrative and denoted by *L* (see Fig. 2A,B). Moreover, both *M* and *R* appear to grow linearly with *L* for the range of narrative lengths we explored, and hence when we plot *R* vs *M*, we also get an approximately linear relationship (see Fig. 2C). This scaling behaviour is very different from what we observed with random lists of words with a characteristic square root scaling, i.e. unsurprisingly, recall of meaningful material is better than for random ones of the same size, even if we discount for better memorization. One of the factors that apparently leads to better recall of narratives is the temporal ordering of recall. When people recall narratives, recall mostly proceeds in the forward direction (see Fig. 3A), probably reflecting the natural order of events in the narrative that cannot be inverted without affecting its coherence. This contrasts with the case of random lists, when recall proceeds in both directions with similar probability (see Fig. 3B), which, according to a model proposed in [Naim et al., 2020] results in the process entering a cycle preventing many words from being recalled.

**Figure 2.**
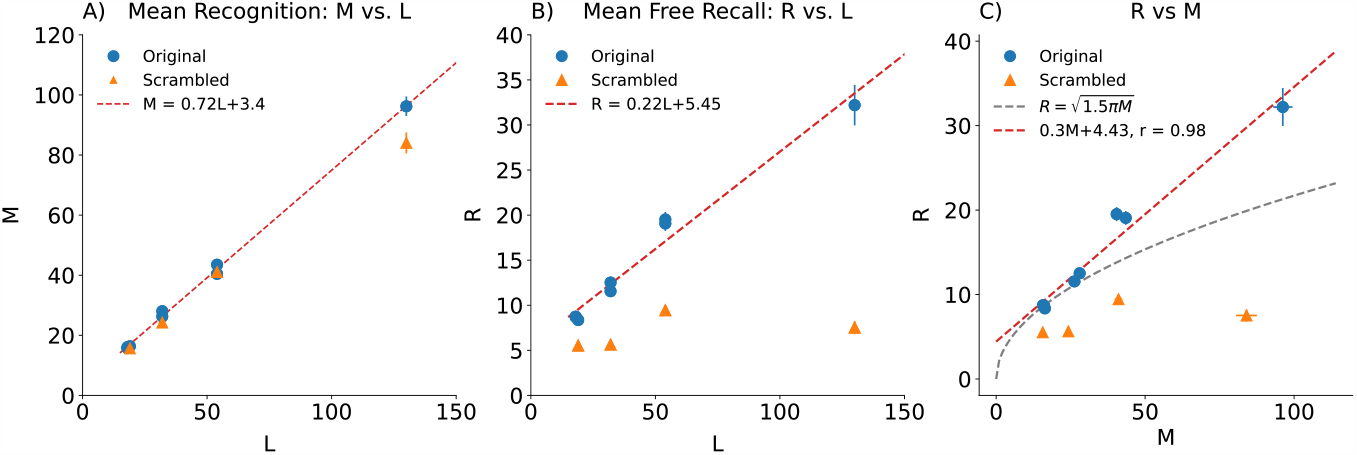
Human performance in recall and recognition experiments for narratives of different length. (A): Estimated number of remembered clauses (M) is plotted as a function of the number of clauses in the narrative (L) measured in recognition experiment. Surprisingly M has similar values in intact and scrambled narrative. (B): Average number of recalled clauses (R) for narratives of different length. In contrast to the M, R drops substantially for scrambled narratives. (C): Average number of recalled clauses vs. number of remembered clauses from the same story. As expected from panels a) and b) the number of retrieved clauses in scrambled narrative is substantially smaller that in intact narrative for the same number of remembered clauses. For comparison we presented the theoretical performance for the random list of words, which describes data well (Naim et al. [2020]). It is clear that there are more clauses recalled in intact narratives than words in lists of random words. Surprisingly, retrieval of scrambled stories is significantly *worse* than random lists, suggesting an active suppression of items in service of generating a coherent recall (participants were implicitly instructed to recall story).

**Figure 3.**
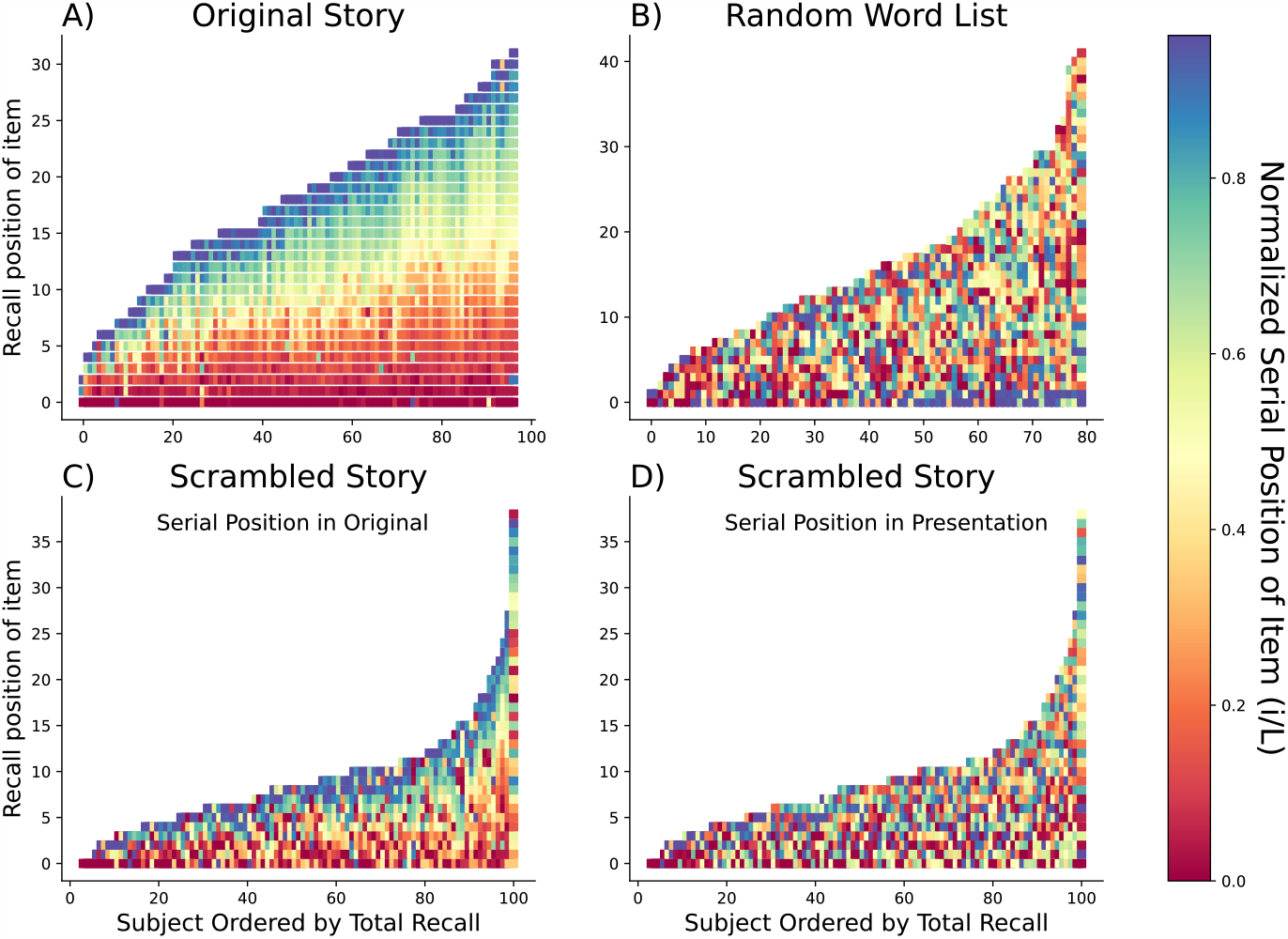
Recall order. Color-coded order of clauses or words for different conditions are shown in all panels. Recalled clauses or words are stacked together vertically (with the first recalled clause at the bottom of a column, and the last recalled clause at the top). The height of the column represents the total number of clauses or words recalled in a given trial. In panels A, B, and D, color code represents serial position of *presentation* of clauses or words, from early (red) to later (blue) in presentation position. Panel (C) is the only exception, in which the color code reflects the serial position of clauses in the *original* (intact) story. (A) shows that recall of coherent stories largely preserves presentation order. (B) recall of random word lists does not preserve presentation order. (C) As with random lists, the recall of a scrambled story does not preserve presentation order, but rather appear to reconstruct the original order of the story, as seen from the color gradients in panel (B). Apparently, random words and scrambled stories are recalled in random order considering their presentation order, but people perform some unscrambling of the scrambled stories as can be seen in (C) - there is tendency of recalled clauses being in the order of original unscrambled narrative. The participants construct a mental representation of the scrambled narrative which is evidently close to its original form. Recall consequently does not reflect input sequence, but rather the original sequence of the clauses.

### 2.3 Meaning and memory

As we mentioned above, people’s recall is strongly influenced by narrative comprehension, such that clauses that are most important in communicating the summary of the narrative are the ones that are recalled by most of the participants. We found however that recognition is not so strongly affected by meaning. This can be observed by evenly dividing all the clauses used in our experiments into subsequent bins according to their recall probabilities, and calculating the average recognition performance for all the clauses in each bin. Surprisingly, there is very little increase of recognition with recall probability across the clauses, such that clauses with highest and lowest average *P*_*rec*_ only differ in their *P*_*h*_ by less than 0.15 (see Fig. 4; c.f. [Thorndyke and Yekovich, 1980; Yekovich and Thorndyke, 1981]).

**Figure 4.**
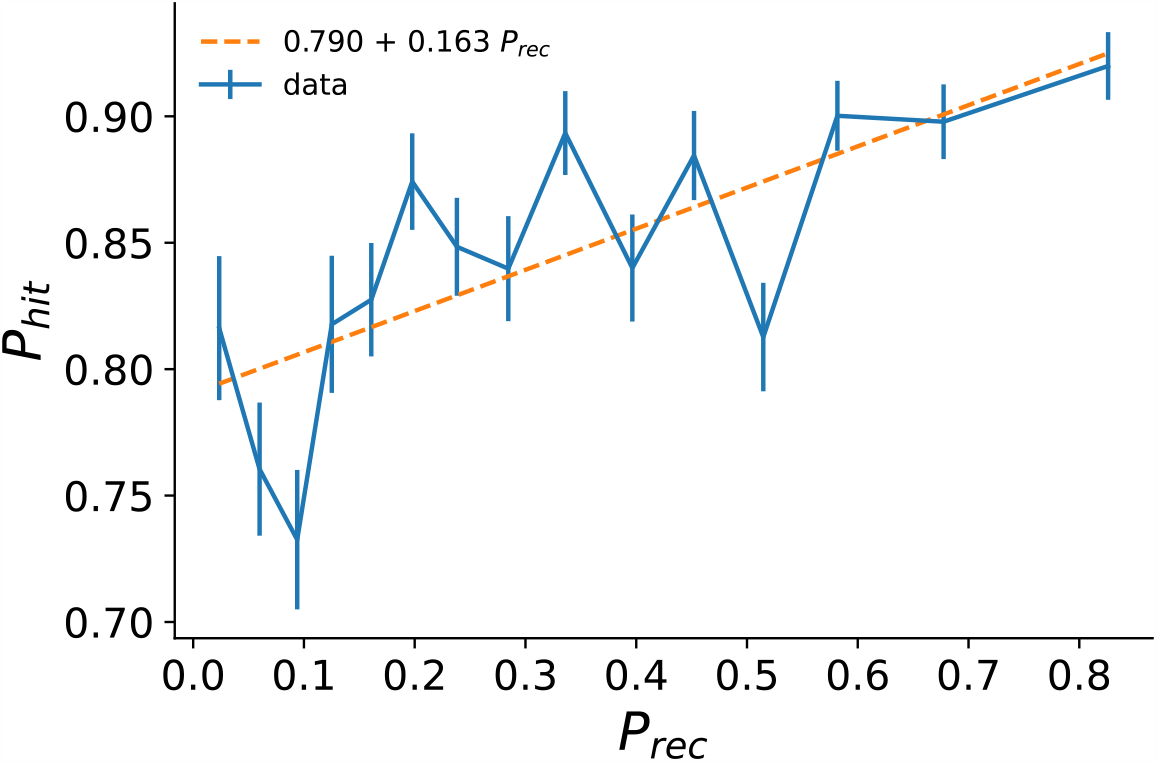
Recognition vs recall performance across different clauses. Clauses from all the narratives used in this study were divided evenly into 15 bins according to their *P*_*rec*_, and the average *P*_*h*_ for the clauses in each bin was computed and plotted against the center of the corresponding bin.

To further elucidate the role of meaning in memory, we repeated our experiments with another group of participants after randomly scrambling the order of clauses, thus making comprehension much more difficult if not impossible. We found that, unsurprisingly, recall of scrambled narratives is much poorer then the original ones (Fig. 2B,C). Recognition performance for scrambled narratives however is practically the same (Fig. 2A). This result indicates that memory encoding of clauses is not significantly affected by the structure and meaning of the narrative. Interestingly, the order in which people tend to recall clauses from a scrambled narrative corresponds much better to the order of these clauses in the original narrative than in the presented, scrambled one (see Fig. 3C,D), indicating that even in this situation people are trying to comprehend the meaning of the narrative rather than processing the input as a random list of unconnected clauses. This might explain why recall of scrambled narratives appears to be *worse* than recall of random word lists of the same size.

## 3 Previous Work

The experimental study of memory for narratives can be traced back to the highly influential descriptive work of Bartlett [1932]. This and followup work introduced the idea that the encoding of memory for narratives is a process of abstraction [Gomulicki, 1956], and that subsequent recall is in large part a generative process driven by a participant’s prior knowledge and biases. This line of thought was formalized much later in theories of narrative structure involving schemas, scripts, frames, and story grammars [Alba and Hasher, 1983; Rumelhart, 1975]. By now, there is ample support for an abstracting process for memory encoding, and the existence of schematic structures which guide recall [Baldassano et al., 2018].

A parallel line of research into narrative structure originated in the field of sociolinguistics by Labov and Waletzky [1966]. In carrying out linguistic fieldwork to analyze spoken dialects of English, the authors found that personal narratives of emotionally charged events tented to elicit the best examples of local dialect. Furthermore, the resulting narratives, which were produced spontaneously and by non-expert storytellers, tended to be very regular in their structural properties. Of particular significance to us was the observation that stories were typically told in the order in which they experienced by the speaker, which Labov encapsulates in his “egocentric principle” and “no flashback” constraint Labov [2013]. This lends some support to the strong iconicity assumption [Zwaan, 1996], which states that readers (or listeners) will interpret the order in which events are reported as reflecting the actual chronological order in which they occurred.

However, serial order alone likely cannot explain the rich structure we observe in narrative recall. Indeed, temporal order provides just a single axis along which events in a narrative are organized. In addition to simple serial order, events or clauses can have causal relations (e.g. A causes B) Trabasso and van den Broek [1985]; Trabasso and Sperry [1985]; Lee and Chen [2022], inferential relations (e.g. A implies B), and superordinate or constituency relations (e.g. A consists of B) Black and Bower [1979]. These relations conspire to give narratives hierarchical structure [Kintsch, 1998]. Story grammars provide one natural mechanism whereby the hierarchical structure arises, and served as early inspiration for studying structure dependence in narrative recall and recognition.

In this direction, Yekovich and Thorndyke [1981]; Thorndyke and Yekovich [1980] performed recall and recognition experiments to test how encoding and processing depend on hierarchical structure. Their recognition experiments included only old, paraphrase, and false statements in the test phase. The false statements tested were appropriate (invoking the appropriate agents or actions), but inconsistent with or contradictory to the story. Similar to our results, they observe that recognition is uncorrelated with recall. Furthermore, whereas recall was argued to be sensitive to the structure of a narrative, recognition appeared to have no such sensitivity.

In another experiment seeking to isolate the structure sensitivity of narrative processing, Kintsch et al. [1977] tested processing of scrambled narratives, in which paragraphs of a text were presented to participants in random order. Participants were able to produce coherent summaries of such scrambled text, which were indistinguishable from summaries produced by participants reading the coherent text. It is possible that scrambling larger units (paragraphs in this case, versus clauses in our experiments) produced an overall more comprehensible text, making descrambling easier. Nevertheless, we still observe descrambling, but of lower quality (see Figure 3C).

Scaling laws for memory have been observed for random unstructured lists (pictures, words, etc) [Standing, 1973; Murray et al., 1976; Naim et al., 2020]. Two important takeaways from these works are the following: memory typically exhibits power-law scaling with list length, and the retrieval process appears to be universal. Surprisingly, there is very little work which considers the scaling of narrative memory with story length, let alone attempts to quantify it. The only work we are aware of is [Glenn, 1978], which measured average recall as a function of story lengths up to 83 “informational units”, which include clauses as well as noun and adjective phrases. The experimental design was motivated largely by questions about story grammars, and therefore the episodic structure of the narratives (in the story grammar sense developed by the authors) was kept constant while descriptive phrases and clauses (so called ‘informational units’) were added to increase the length of the story. Therefore, the added statements are arguably a kind of filler, not contributing any significant additional meaning or structure to the underlying narrative. This might account for the seemingly sublinear scaling of their mean recall with story length, compared to our linear results (Figure 2). There are other significant differences, including the test population (24 second-grade schoolchildren), the stimulus input format (spoken narratives), and the recall format (spoken recall).

The choice of “informational units” in [Glenn, 1978] also differs from our choice of clauses, and is similar to segmentations that use propositional analysis [Kintsch and van Dijk, 1978], or pausal breaks [Johnson, 1970]. An important feature of these different units is that they provide a more fine-grained segmentation of a narrative - a clause can consist of many propositions or pausal breaks, but not the other way around. However, these smaller units would not have a narrative function in the Labovian sense.

## 4 Discussion

In this contribution, we describe a new way to study human memory for meaningful narratives with the help of large language models. Together with using internet platforms for performing experiments, this technology enables a qualitative leap in the amount of data one can collect and analyze. In particular, we describe the prompts that we used to make GPT-4 generate new narratives of a particular type and size, score multiple human recalls of these narratives by identifying which clauses were recalled, and generating plausible lures for recognition experiments. Having a large amount of data is important for memory research because, as opposed to e.g. sensory processing that is believed to be largely uniform across people [Read, 2015], the way people remember and recall meaningful material is highly variable. Hence, only through statistical analysis can some general patterns be uncovered. In particular, we considered how recognition and recall performance scales up for narratives of increasing length. We found that approximately the same fraction of narrative clauses are recognized and recalled when narratives become longer, in the range of 20 to 130 clauses that we considered for this analysis. We expect that as narratives become longer, this trend will not persist because people will start summarising larger and larger narrative chunks into single sentences, i.e. simply counting the number of recalled clauses will not be a good measure of recall performance and a better recall metric will have to be developed.

We investigated the role of meaning in narrative memory by presenting the participants with the same clauses as in the original narrative but in a scrambled order. These ‘narratives’ are much more difficult, if at all, to make sense of, and indeed their recall was very poor. However, recognition of individual clauses was practically as good as in the original narrative. This surprising observation indicates that the encoding of clauses in memory is not very sensitive to the overall structure of the narrative, and only in the process of recall does the meaning of the narrative play a major role. This finding is consistent with the observations of [Thorndyke and Yekovich, 1980; Yekovich and Thorndyke, 1981]. It also provides a striking confirmation of the generally held wisdom that while recall is structure sensitive, recognition need not be [Kintsch, 1998]. It is still possible, however, that these trends will change when longer narratives are considered. This will have to be investigated further.

Another interesting observation concerns the fact that as the narrative becomes longer, the range of recall probabilities for different clauses remains very wide, e.g. there are always some clauses that are recalled by most of the participants and others that are almost never recalled. In contrast, the probability to recall words from a random list of words decreases with the length of the list, with the exception of the last few words (due to the recency effect) [Murdock Jr, 1960]. It would be an interesting theoretical direction for future studies to try to identify the factors that could predict how well a given clause will be recalled in a given narrative.

We focused on first-person spoken stories. These were personal accounts of important events, shared naturally and informally. This way, they lacked the refinement of crafted stories, which may utilize tricks to improve memorability. It would be interesting to see how scaling of memory is affected by such expertly told or literary stories. Evidently there is an impact, considering that many stories in the oral tradition endure over very long timescales [Rubin, 1995; Nunn, 2018]. A striking example of this in a more controlled lab setting had participants trained to construct a narrative in which they embedded a random list of words [Bower and Clark, 1969]. This work found that employing such a mnemonic improved recall to nearly perfect for up to 12 consecutively learned lists.

While practice and rehearsal are necessary for preserving narratives in oral tradition, our results suggest that narratives are intrinsically more memorable. We find that memory performance for narratives encountered only once, as measured by the scaling relations in Figure 2, are robustly superior to performance on unstructured lists.

The research conducted for this report relied crucially on a set of LLM input “prompts”, i,e, instructions, written in standard English, given to the LLM for carrying out various tasks. Roughly speaking, these appear as if they were instructions given to a human research assistant. Quite remarkably, the LLM completes the input string to provide a correct output without any additional training, a phenomenon known as ‘in-context’ learning [Brown et al., 2020]. Since this phenomenon is still not fully understood, we had to resort to a good amount of trial-and-error and fine-tuning in designing the prompts used in our analysis. We provide all of the prompts used in our experiments in appendix A. The specific model we utilize in most of the paper is OpenAI’s GPT-4. However, while we believe the capabilities necessary to carry out our experiments are not limited to this model, it is an open question whether the prompts we use can be transferred to different models.

In summary, using LLMs in conjunction with internet platforms for performing experiments is a powerful new tool that could bring significant new advances in understanding human memory.

## 5 Methods

With the aim of conducting a large scale study on memory for natural continuous material (personal narratives in this case) we required an automated procedure that would facilitate measuring human recall memory performance, since manual scoring of recalls is very labor-intensive and thus limits the ability to analyse large datasets. We were able to achieve this through the use of Large Language Models (LLMs) and we assessed the reliability of our pipeline by comparing it to human scoring performed by the authors. Our dataset was generated by conducting recall and recognition experiments online, recruiting participants through a crowd-sourcing platform. All segments of this study are detailed below.

### Stimulus Set - Narrative pool

Nearly all of the stimuli we use are generated by LLMs and are based on first-person oral narratives taken from socio-linguistic interviews [Labov and Waletzky, 1966; Labov, 2013]^2^. The oral narratives are segmented into clauses in these references, and these are used as templates for the LLM narrative generation. The LLM output is a narrative of equal length (in number of clauses), with very similar narrative-syntactic structure, but involving different subject matters. Two stories were generated from each template for lengths L = 18, 32, and 54. Two additional narratives were directly taken from Refs. [Labov and Waletzky, 1966; Labov, 2013], one with L = 19 (‘boyscout’ Story 10), and the other with L = 130 which was minimally edited to remove punctuation due to speech breaks, in order to increase readability (‘stein’ Story 9). More details of the narrative generation by LLMs can be found in appendix A.1, along with a sample narrative template in Argument 1 and examples of generated outputs in Completion 1. All narratives used as stimuli can be found in Appendix F.

For the purpose of evaluating the reliability of recall scoring of LLMs and their similarity to human scoring, we generated and segmented a different narrative based on the ‘boyscout’ story (Story 10, L=19). This stage began before the rollout of `gpt-4` and for this reason we document the evaluation process separately in Appendix H. The narrative generation step in this part produced variable length narratives in prose, which we had to subsequently segment using GPT-3. As a result, this narrative generation procedure did not keep the same number of clauses as the story it was based on (‘boyscout’).

### Experimental Design

Participants were recruited online through the ‘Prolific’ platform (www.prolific.com) and experiments were conducted on participant’s browser. Only candidates that indicated English as their first language were allowed to participate. Participants were initially greeted with a short description of the experiment and an informed consent form. The trial was initiated by a button press. After a three second counter, a narrative was presented in the form of rolling text (marquee) in black font in the middle of a white screen. All narratives were animated in constant speed with a total duration calculated as the character span of the story divided by 250 (resulting in a character moving with a speed of approximately 25 characters/s) while the font size has been set to 40px. Once the marquee for the narrative has traversed outside the screen (all characters shown and disappeared), the testing phase was triggered automatically. This marquee style presentation was chosen because it allowed for comfortable reading while fixing the presentation duration for all participants and simultaneously preventing revisiting of already read material.

In the free recall experiments, the testing phase consisted of a textbox and a prompt to recall the story as close as possible to the original one. Once participants finished typing their recall of the narrative, they submitted their response with a button press and the experiment was concluded.

For the testing phase of the recognition experiment, 10 queries were sequentially presented. In each query, the participant was shown a single clause at random, either from the just presented narrative (old) or a lure (new). They were tasked to select whether they remembered seeing the presented clause by pressing “Yes” if they thought the clause appeared in the narrative and “No” otherwise. We did not observe any obvious signatures of output interference [Criss et al., 2011] (see Appendix C) and therefore used all queries in the subsequent analysis. Lures were generated by asking the LLM to take a given narrative segmentation, and insert novel clauses between each existing clause. This ensures that the lures are distinct from the true clauses, but still fit within the overall context of the story. For instance, this avoids lures which might mention “dolphins” if the story is about boy scouts. The prompt used to generate lures and an example completion by `gpt-4` are given in Appendix A.2.

### Analysis

Analysis was conducted through custom Python scripts. For recognition memory, in order to estimate total encoded memory *M* from Equation (1), we used population and test trial averaged hit rate (true positive probability *P*_*h*_) and false alarm rate (false positive probability *P*_*f*_). Standard error was computed using statistical bootstrap [Efron and Tibshirani, 1994].

Recall scoring was done using the OpenAI model `gpt-4-0613` (a GPT-4 model which receives no updates) based on the clause segmentation of the narratives. For each participant’s recall, `gpt-4-0613` was instructed to loop through each clause of the given narrative (as presented) and examine whether the information that this clause provided was present in some form in the participant’s recall and the corresponding passage. The numbers of all clauses evaluated as being recalled were given at the end of the output in the form of a list. The full prompt we use for scoring recalls is given in Prompt 3, which takes three arguments: the narrative stimulus in prose (e.g. Argument 2), the numbered clause segmentation of the narrative stimulus (e.g. Argument 3), and the participant’s recall (e.g. Argument 4). A sample completion is provided in Completion 3.

Separately, to evaluate the similarity of recall scoring between humans and LLMs, 3 authors performed manual scoring of 30 recalls of Argument 5 using the same procedure, evaluating whether each clause was present in the recall or not.

### Random List of Nouns Experiment

We performed an additional experiment with a list of 32 nouns (see Appendix G) that were randomly selected from the pool of nouns used in [Naim et al., 2020]. The experimental protocol was exactly the same as in that work with presentation speed 1.5 sec per word. 105 participants were recruited using the Prolific online platform, with each participant accepting informed consent prior to the beginning of a trial.

## Data And Code Availability

Experimental data as well as code for the online experiments and the analysis presented above can be found in https://github.com/mkatkov/llm-narrative-analysis.

## Acknowledgements

TC acknowledges the support of the Eric and Wendy Schmidt Membership in Biology, the Simons Foundation, and the Starr Foundation Member Fund in Biology at the Institute for Advanced Study. MT is supported by the Foundation Adelis and the Simons Foundation. We thank Omri Barak, Danqi Chen, Michael Douglas, Yadin Dudai, and Ariel Goldstein for helpful conversations. We thank Stefanos Gkouveris for helpful comments on Javascript development.

## Appendix A. Prompts and Completions

A significant part of the successful use of LLMs is the effective design of inputs or “prompts”, which amount to instructions, written in standard English, for carrying out a particular task.

More precisely, the input to a LLM is a string, which we interchangeably refer to as the prompt or the context. The output of the LLM is also a string. This is also referred to as a “completion”, since LLMs are trained to complete text fragments given to them as inputs.

Despite the accumulation of wisdom on prompt engineering, we still resorted to a good amount of trial-and-error and fine-tuning in designing the prompts used in our analysis. Since these prompts are tantamount to algorithms written for LLMs, we present them below in pseudocode boxes. In the following sections, we exhaustively detail the prompts used, along with examples of LLM outputs (i.e. completions). The specific model we utilize in most of the paper is OpenAI’s GPT-4. For the data analysis, in the interest of reproducibility, we opted to use the deprecated model `gpt-4-0613` which does not receive updates. In appendix B.1, we show how scoring using the latest model `gpt-4` can change with time as a result of OpenAI’s regular model updating.

### A.1 Narrative Generation

We started with a template narrative, selected from the collection of oral narratives in [Labov, 2013], and instructed the LLM to produce variations of the story which changed the surface form, but kept the overall structure (e.g. number of clauses). For these stories, we used the segmentation given by Labov [2013], wherein the narratives are segmented word-for-word into linguistic clauses. Here is a sample of the prompt, with arguments shown presented in a box enclosed in brackets, like `{ this }`.

#### Prompt 1

**Narrative Generation**

~~~
This is a true personal narrative about a single event in someone’s life. It has exactly { N } clauses:
{ template narrative }
Generate a new personal narrative that is unique and about something completely different.
Try to keep the overall narrative structure of the personal narrative
above, but change as much of the subject matter and action as possible. Do
not just use the narrative and replace key persons, places and things. Make it
completely new. This new narrative must also contain exactly { N } clauses.
~~~

As an example, we used the story of Jacob Schissel from [Labov, 2013] as a template narrative which has `N` = 18:

#### Argument 1

**Example template narrative**

~~~
1. My brother put a knife in my head.
2. This was just a few days after my father had died
3. and we were sitting shiva.
4. And the reason the fight started,
5. he saw a rat out in the yard or
6. – this was out in Coney Island –
7. and he started talk about it.
8. And my mother had just sat down to have a cup of coffee
9. and I told him to cut it out.
10. ’Course kids, y’know, he don’t hafta listen to me.
11. So that’s when I grabbed his arm
12. and twisted it up behind him.
13. When I let go his arm, there was a knife on the table,
14. he just picked it up
15. and he let me have it.
16. And …; I started bleeding – like a pig.
17. And naturally first thing to do, run to the doctor,
18. and the doctor just says, “Just about this much more,” he says, “and you’d a been dead.
~~~

The output of the LLM upon processing this prompt was a story which clearly bore a structural relation to the original, but contained different agents involved in a completely different scenario. We provide an example completion below obtained using gpt-4 with temperature T = 0.6. This example was used in experiments, and also appears in Story 3 next to another generated narrative (Story 4).

#### Completion 1 from Prompt 1, Argument 1

~~~
1. My best friend pushed me into the pool.
2. This was right after I had broken up with my boyfriend
3. and I was feeling really down.
4. The reason she did it,
5. she thought I needed a distraction or
6. - this was at her house in the suburbs -
7. and she wanted to cheer me up.
8. I had just sat down with a plate of snacks
9. and I told her I was fine.
10. ’Course friends, y’know, they don’t always believe you.
11. So that’s when she sneaked up behind me
12. and gave me a little shove.
13. When I surfaced from the water, there was a float nearby,
14. she just grabbed it
15. and tossed it to me.
16. And …; I started laughing - uncontrollably.
17. And naturally the first thing to do, hug my friend,
18. and I just said, “Just a little more,” I said, “and I would’ve been drowning in my sorrows.
~~~

The resulting narratives generated from this procedure vary in their coherence. For our experiments, we chose the most coherent generated narratives

The resulting narratives generated from this procedure vary in their coherence. For our experiments, we chose the most coherent generated narratives.

### A.2 Lure Generation

We utilized `gpt-4` at *T* = 0.3 with the following prompt:

#### Prompt 2

**Lure Generation**

~~~
{ segmentation }
The items above all fit together to tell a story. Add more items of roughly
the same length, numbered 1.5, 2.5, and so on, interleaving the existing items,
elaborating on the story, and without repetition. These new items should introduce
completely new plot elements, but still make sense in the context of the rest of
the story. Add as many items as possible.
~~~

The following is an example completion using the template narrative segmentation (Argument 1). For purposes of presentation, we present the lures along with the original clauses from the narrative. The completion does *not* include the original clauses, which appear in gray, and only consists of the text shown in black.

#### Completion 2 from Prompt 2

~~~
1. My brother put a knife in my head.
1.5. It was a tense time for our family, with everyone on edge.
2. This was just a few days after my father had died
2.5. We were all grieving in our own ways, trying to cope with the loss.
3. and we were sitting shiva.
3.5. Our house was filled with relatives and friends, offering their condolences.
4. And the reason the fight started,
4.5. My brother had always been a bit of a troublemaker, but I never thought he’d go this far.
5. he saw a rat out in the yard or
5.5. He was always fascinated by the creatures that roamed the streets and alleys.
6. – this was out in Coney Island –
6.5. The sun was setting, casting an eerie glow on the scene.
7. and he started talk about it.
7.5. He started describing the rat in gruesome detail, trying to get a rise out of us.
8. And my mother had just sat down to have a cup of coffee
8.5. She looked so tired and worn out, I just wanted her to have a moment of peace.
9. and I told him to cut it out.
9.5. I tried to keep my voice calm, but I could feel my anger rising.
10. ’Course kids, y’know, he don’t hafta listen to me.
10.5. He just smirked at me, clearly enjoying the attention.
11. So that’s when I grabbed his arm
11.5. I didn’t want to hurt him, just get him to stop his nonsense.
12. and twisted it up behind him.
12.5. He let out a yelp of pain, but I didn’t think I’d done any real damage.
13. When I let go his arm, there was a knife on the table,
13.5. I didn’t even see him reach for it, he was so quick.
14. he just picked it up
14.5. I barely had time to react before the blade was coming towards me.
15. and he let me have it.
15.5. The pain was sharp and intense, like nothing I’d ever felt before.
16. And …; I started bleeding – like a pig.
16.5. I could feel the warmth of my blood soaking through my clothes.
17. And naturally first thing to do, run to the doctor,
17.5. I stumbled out of the house, desperate for help.
18. and the doctor just says, “Just about this much more,” he says, “and you’d a been dead.
18.5. I couldn’t help but think how close I’d come to joining my father.
~~~

### A.3 Recall Scoring

In order to score recall, we utilize zero-shot prompting with OpenAI’s deprecated model `gpt-4-0613`. We chose to use this particular model for the benefit of reproducibility, since we have found that results from the following analysis can drift in time as OpenAI updates its model (see Appendix B.1).

The recall scoring Prompt 3, shown below, was a block of text, represented as a python string, which took as arguments the original narrative, the numbered segmentation of the narrative, and a single participant’s recall of the narrative. The prompt was constructed so that it would identify which clauses from the narrative are present in the recall. A consequence of this scoring procedure is that a single clause in the recall can encode multiple clauses from the original narrative.`gpt-4` appeared to have no trouble identifying such compressed clauses in the recall, and frequently would identify multiple original clauses within a single clause in the participant’s recall. An example of this can be seen in Completion 3, where the same passage in the recall (“even the scoutmaster looked on”) is counted as recalling both clauses 15 and 16 from the original narrative (Argument 2).

Below, we provide the scoring prompt, as well as some example arguments, and the resulting LLM completion.

#### Prompt 3

**Recall Scoring**

~~~
This is the original text:
{ narrative }
It can be broken down into the following independent pieces of information:
{ segmentation }
Here is an alternative version of the original text where some of the above pieces
of information may be missing:
{ recall }
For each of the numbered information pieces of the list above, evaluate whether
the information of each piece is given in the alternative version of the story,
stating the number and showing the corresponding passage from the alternative story
it is given in. After, write all the numbers of the pieces that are given in the
alternative version of the story in a set of brackets at the end of the response.
~~~

#### Argument 2

**Example narrative**

~~~
Yeah, I was in the boy scouts at the time. And we was doing the 50-yard dash,
racing, but we was at the pier, marked off and so we was doing the 50-yard dash.
There was about 8 or 9 of us, you know, going down, coming back. And going down
the third time, I caught cramps and I started yelling “Help!”, but the fellows
didn’t believe me, you know. They thought I was just trying to catch up, because I
was going on or slowing down. So all of them kept going. They leave me. And so I
started going down. Scoutmaster was up there. He was watching me. But he didn’t
pay me no attention either. And for no reason at all there was another guy, who
had just walked up that minute… He just jumped over and grabbed me.
~~~

#### Argument 3

**Example segmentation**

~~~
1. Yeah, I was in the boy scouts at the time.
2. And we was doing the 50-yard dash,
3. racing,
4. but we was at the pier, marked off,
5. and so we was doing the 50-yard dash.
6. There was about 8 or 9 of us, you know, going down, coming back.
7. And, going down the third time, I caught cramps
8. and I started yelling “Help!”,
9. but the fellows didn’t believe me, you know.
10. They thought I was just trying to catch up, because I was going on or slowing down.
11. So all of them kept going.
12. They leave me.
13. And so I started going down.
14. Scoutmaster was up there.
15. He was watching me.
16. But he didn’t pay me no attention either.
17. And for no reason at all there was another guy, who had just walked up that minute…
18. He just jumped over
19. and grabbed me.
~~~

#### Argument 4

**Example recall**

~~~
A boy was with the scoutmaster and he was going up and down and on the third time
of going up the pier he had cramps and shouted ‘help’ but no-one believed him, even
the scoutmaster looked on by until this guy came out of nowhere and grabbed the boy
and saved him.
~~~

#### Completion 3 from Prompt 3

~~~
1. Not given
2. Not given
3. Not given
4. Not given
5. Not given
6. Not given
7. Given - “on the third time of going up the pier he had cramps”
8. Given - “and shouted ‘help’”
9. Given - “but no-one believed him”
10. Not given
11. Not given
12. Not given
13. Not given
14. Given - “was with the scoutmaster”
15. Given - “even the scoutmaster looked on”
16. Given - “even the scoutmaster looked on”
17. Given - “until this guy came out of nowhere”
18. Not given
19. Given - “and grabbed the boy and saved him”
(7, 8, 9, 14, 15, 16, 17, 19)
~~~

We note here that the completion does not always appear as in this example. Some minor deviations might occur (e.g. parentheses replaced with brackets), or the format of the answers change (e.g. instead of “Not Given”, some completions give “There is no corresponding passage for “).

### A.4 Ordered Recall Scoring

In order to extract the order in which the original clauses appeared in a participant’s recall, we added an extra step to the recall scoring discussed above. For a given narrative, segmentation, and recall, we concatenated Prompt 3 with its Completion 3, and appended to this the following additional prompt:

#### Prompt 4

**Ordered Recall Scoring**

~~~
Now repeat the alternative version of the narrative with the number of the
independent piece of information inserted next to the location in which it appears
in the alternative version. Then, list the numbers separately in the order in
which they appear in the alternative story immediately above. The final list of
numbers should be enclosed in parentheses.
~~~

Therefore, the total input or prompt for ordered recall scoring would be Prompt 3 + Completion 3 + Prompt 4. The output for this example was the following:

#### Completion 4

**from Prompt 3 + Completion 3 + Prompt 4**

~~~
A boy (14) was with the scoutmaster and he was going up and down and on the third
time of going up the pier he had cramps (7) and shouted ‘help’ (8) but no-one
believed him (9), even the scoutmaster looked on (15, 16) by until this guy came
out of nowhere (17) and grabbed the boy and saved him (19).
(14, 7, 8, 9, 15, 16, 17, 19)
~~~

## Appendix B. Reliability of LLM scoring

Here we report additional details of the reliability for LLM narrative scoring.

In scoring the recalls, the human scorers (who are the authors on this paper) did not develop overly sophisticate scoring protocols. The directions were intuitive, essentially boiling down to “did this clause from the original story appear in the recalled story”. Despite the ambiguity of this procedure, there was very high inter-scorer correlation.

### B.1 Drift in Performance over time

The models offered through the OpenAI API are known to undergo regular updates. We found that within the span of two months, the correlation coefficient with mean human performance rose by 0.05. We compare `gpt-4` called via OpenAI’s API on 05/23/23, the same model `gpt-4` called less than two months later on 07/03/23, with the deprecated model `gpt-4-0613` (called after 06/13/23). Fortunately, the frozen model `gpt-4-0613`, which we use throughout the paper, is essentially as good as the most recent version.

### B.2 Comparison between different LLMs

We compared different LLMs available through the OpenAI API. In order of increasing number of learned parameters (model size), these were GPT-3 (`text-davinci-003`), GPT-3.5 (or ChatGPT, `gpt-3.5-turbo`) and GPT-4 (`gpt-4-0613`). The same scoring Prompt 3 was used for all of these models. The results are visualized in Figure 5. We find that GPT-4 is qualitatively better than the smaller models (compare correlation between human and LLM scoring in Figure 5B). Furthermore, increasing size does not seem to be sufficient to increase performance, as illustrated by the severe drop in performance by GPT-3.5. As seen in Figure 5(A,B), GPT-3.5 appears to score more generously than the other models, resulting in a systematic upward bias of nearly all recall probabilities. It is possible that this reflects the sensitivity of these models to prompts, and that with appropriate tuning of the scoring prompt, the performance of GPT-3.5 could be improved.

**Figure 5.**
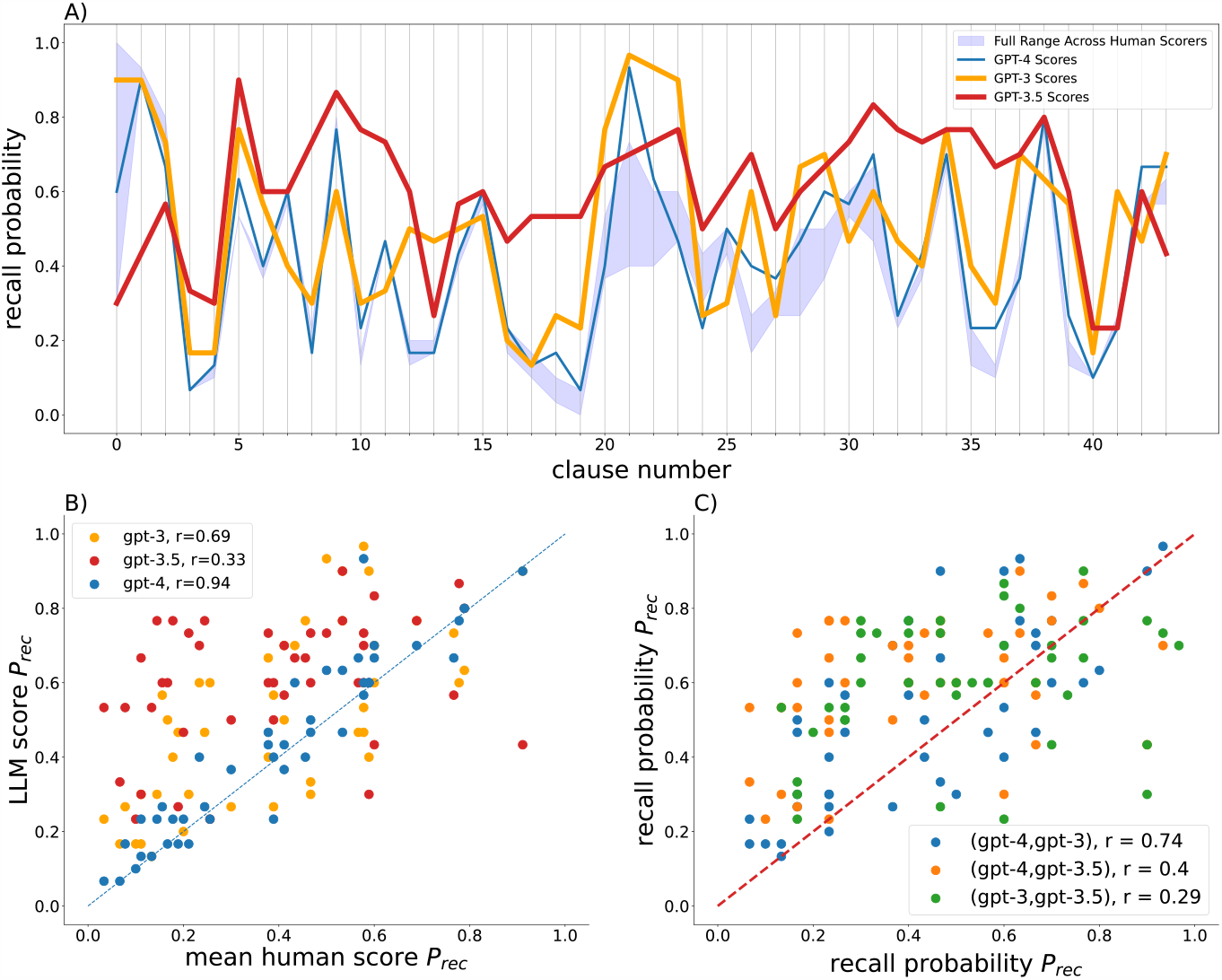
Reliability of different LLMs for scoring narratives. We observed that performance on human recall scoring was not monotonic in the size of the LLM. (A) shows *P*_*rec*_ for each clause as computed by LLMs of three different sizes, in order of increasing size: GPT-3 (OpenAI’s API model `text-davinci-003`), GPT-3.5 (or ChatGPT, model `gpt-3.5-turbo`), and GPT-4 (`gpt-4-0613`). GPT-3.5 scores appear systematically biased to be higher than the mean human scores, whereas GPT-3 roughly follows the human trend. (B) shows correlations between these models and mean human performance. (C) shows inter-LLM correlations.

## Appendix C. Lack of Output Interference in Recognition Experiments

Similar to [Criss et al., 2011], we measure the strength of output interference in recognition experiments by tracking the *d*^′^ measure as a function trial number. The discrimination measure used here is conventionally called simply *d*^′^ and is given by the difference in z-score between the hit probability *P*_*h*_ (i.e. true positive rate) and the false alarm probability *P*_*f*_ (i.e. false positive rate). The z-transform is defined

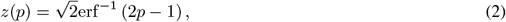

and the discrimination measure is

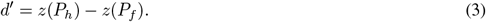

We plot *d*^′^ over the course of the recognition trials in Figure 6 separately for coherent stories and scrambled stories. Output interference is characterized by a *d*^′^ which decreases with trial number, indicating a diminishing ability to discriminate new vs old items.

**Figure 6.**
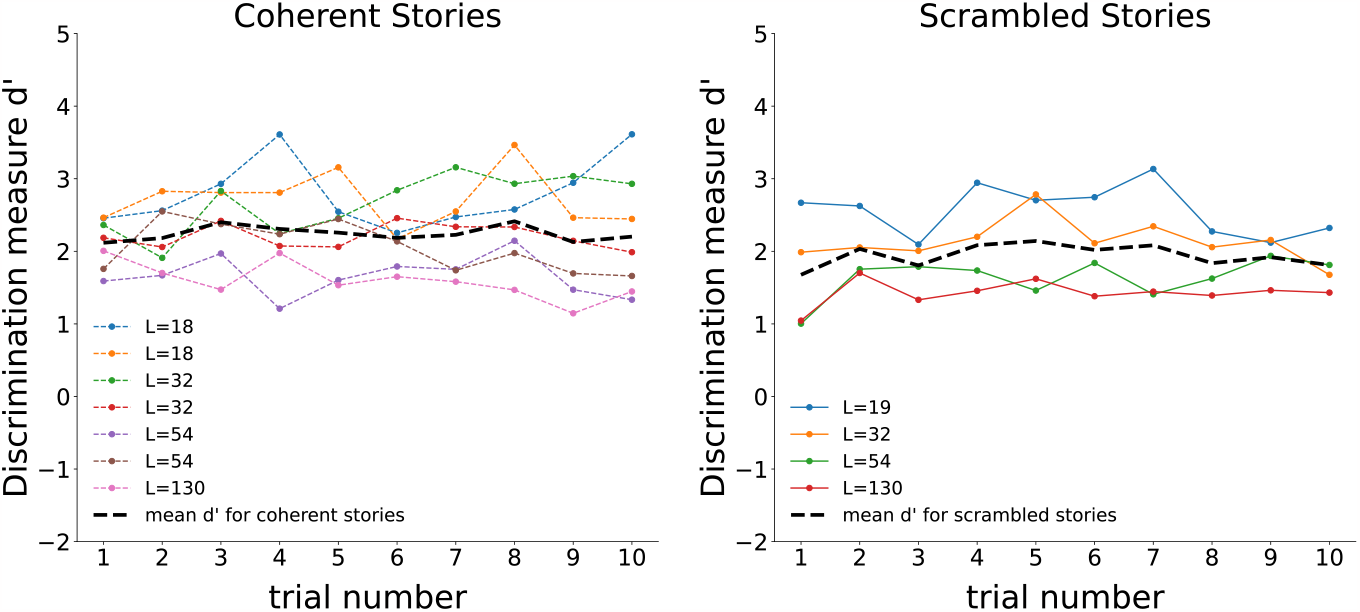
Discrimination measure shows good memory for true clauses which does not change significantly as the recognition trial progresses.

## Appendix D. Recall Probability Curves and Distributions

Here we plot recall probability per clause as a function of the serial position of presentation. We refer to these below as recall probability curves. We compare recall probability curves for narratives to data obtained for random word lists. Figure 7 a shows the recall probability curve for coherent (blue), scrambled (red) stories, alongside the recall probability curve for free recall of a random word list (black dashed). Importantly, for the scrambled recall, the serial position corresponds to the position of *presentation*. Two main points can be gleaned from these figures. First, whereas for random lists there is a pronounced primacy effect, we see no such structure in both coherent and scrambled stories. In fact, highly recalled clauses appear at all positions in the story. Another observation is that the distribution of *P*_*rec*_ for clauses in a narrative covers a much broader range than *P*_*rec*_ for words in a list. This can be seen by looking at the cumulative distribution function of *F* (*p*) = *P* (*P*_*rec*_ *> p*), which is the probability to find *P*_*rec*_ *> p*. As shown in Figure 8A, *F* (*p*) drops to zero around *p* ≈0.6 for random lists, while showing a more gradual decay for coherent narratives. In contrast, Figure 8B shows that recall is significantly impaired for scrambled stories, with *F* (*p*) showing a more precipitous fall to zero similar to random word lists.

**Figure 7.**
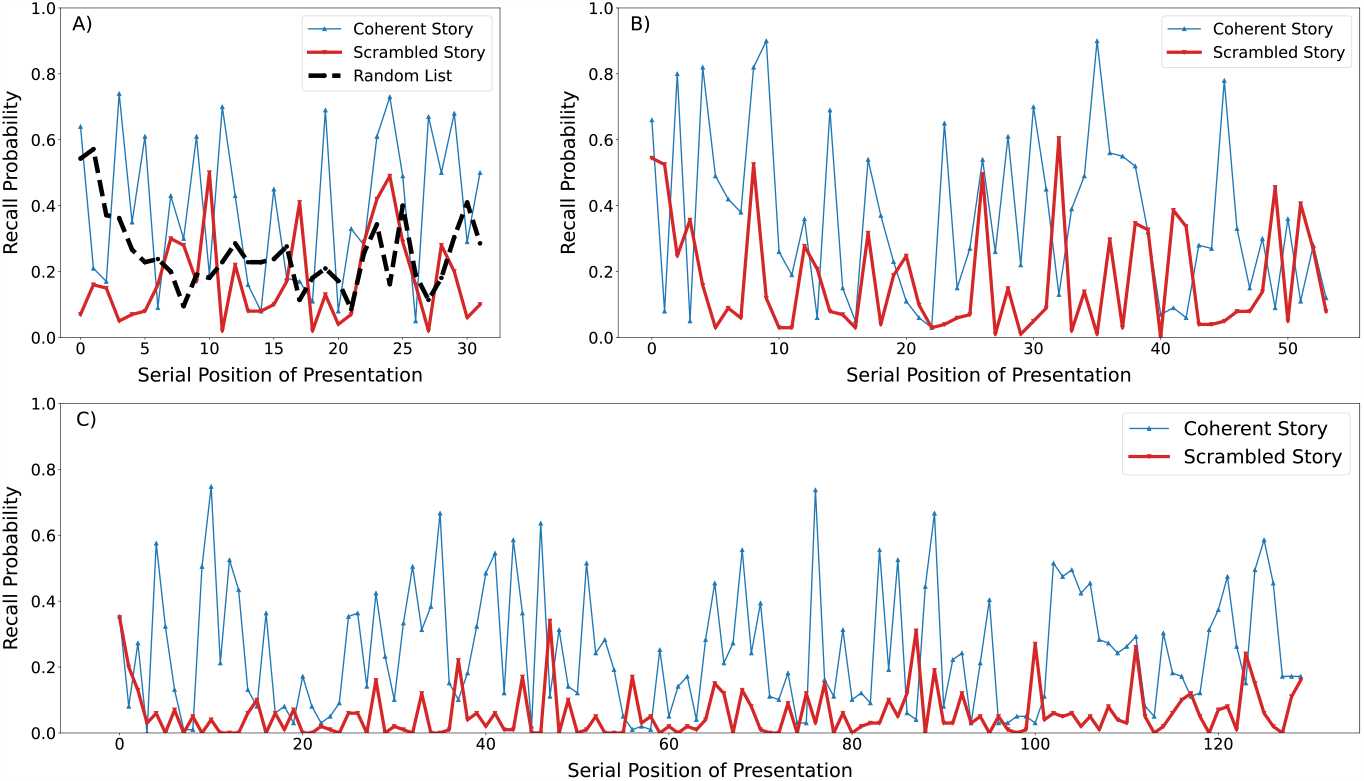
Recall probability per item for narratives and random lists of words. (A) shows the recall probability per clause as a function of the serial position of presentation for a coherent narrative (blue) and the scrambled version of this same narrative (red). As a comparison, we plot the recall probability per *word* as a function of serial position of presentation (black dashed). While the recall probability for the list shows a marked primacy effect, with a general suppression of recall in the bulk of the list, the recall of coherent narratives shows many spikes in *P*_*rec*_ throughout the narrative. And while the scrambled stories have overall much lower recall probabilities, they still show large peaks within the bulk of the presentation. (B) and (C) compare coherent and scrambled narratives of increasing length.

**Figure 8.**
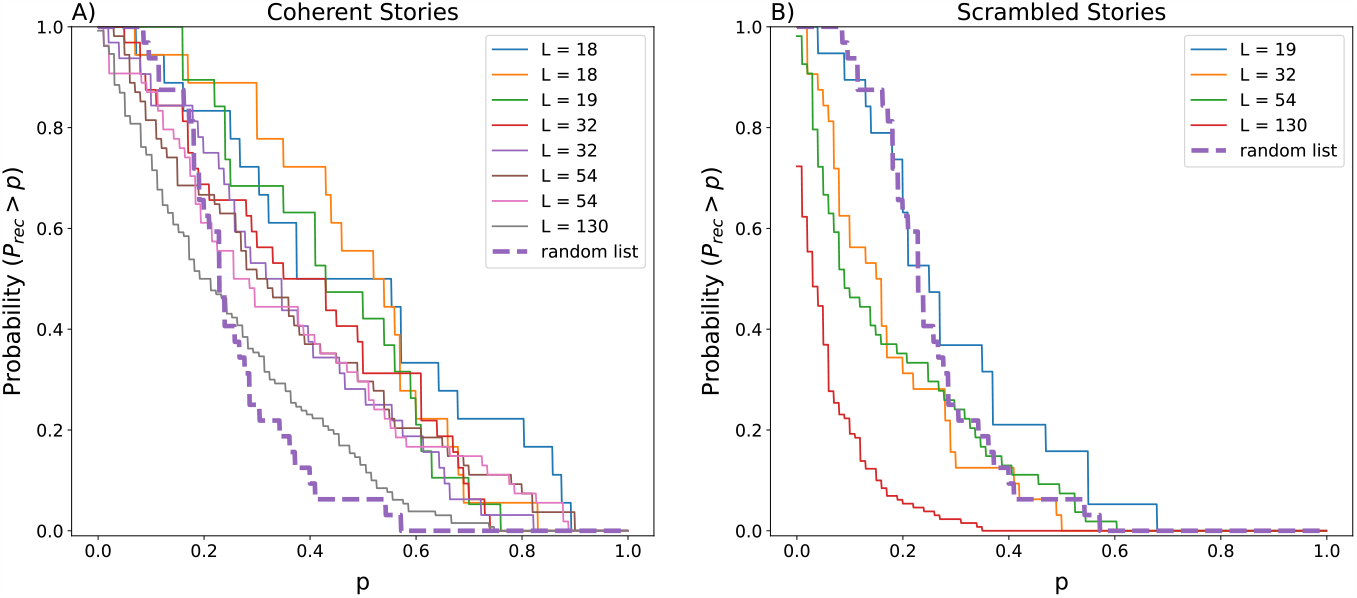
Cumulative distribution of recall probabilities. (A) As compared to random lists (purple dashed), coherent narratives have a slower decline in the cumulative distribution function, indicating a larger number of clauses with a higher recall probability. (B) shows this behavior is reversed for scrambled stories, where the recall probability of clauses appears to uniformly drop, in some cases well below the corresponding distribution for recall of a random list.

However, the recall probability curves for scrambled stories reveals a more subtle effect. While the overall distribution of recall probabilities may be similar to (if not worse than) random lists, there are still spikes in *P*_*rec*_ throughout the bulk of the narrative, suggesting that these texts are not processed as if they were a random list of clauses. In Figure 9, we replot the recall of only the scrambled stories from Figure 7 according the serial position of the clause *in the coherent story*. Below this, we show the correlation between recall probability of a clause appearing in the coherent story versus the scrambled version. While low, the correlations are positive and statistically significant, suggesting that participants are able to identify memorable clauses even in the scrambled scenario, and use this in attempting to construct a coherent recall. This active process of selection might explain why average recall of clauses for scrambled narratives appears *worse* than recall of words in random lists (i.e. why the scrambled narrative data (orange triangles) lie below the square-root scaling (dashed gray) in Figure 2).

**Figure 9.**
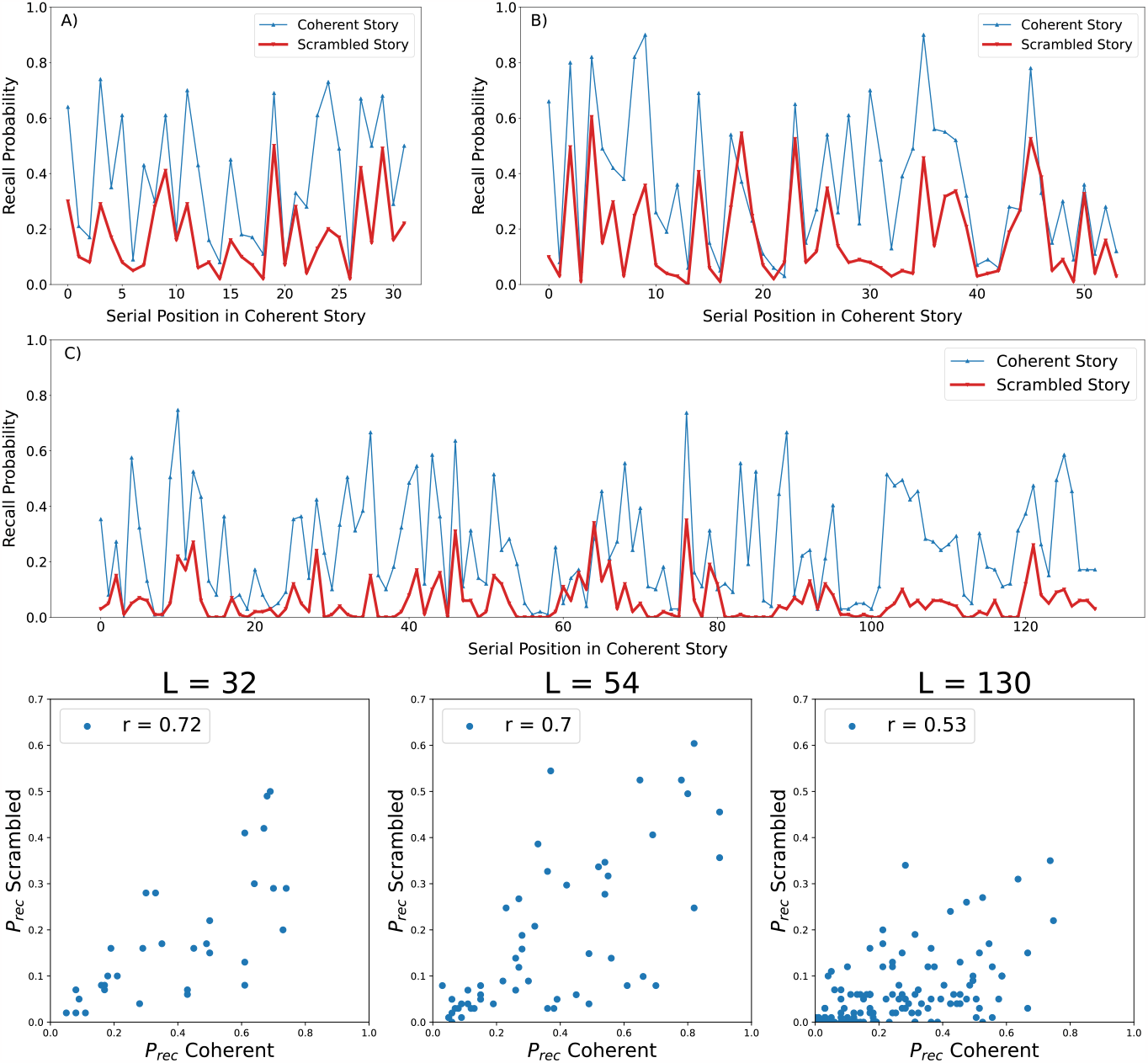
Descrambling of narratives in recall. (A-C) show recall probability as a function of the serial position in the coherent story. Note that for the scrambled *P*_*rec*_ (red curves), this is *not* the presentation order, but rather the original order of the clauses. Plotting the curves in this way, we observe the spikes in recall probability appear to coincide between coherent and scrambled recall. Below, we show correlations between *P*_*rec*_ for a given clause in the coherent and scrambled presentation.

## Appendix E. Clauses with High Recall Probability

We noted that clauses recalled with higher probability tend to encapsulate a summarized version of the presented narrative. While not all nuances are retained, they collectively outline the principal storyline. Presented below are examples from stories of varying lengths. It’s important to note that we present the recalled clauses exactly as they appeared in the original story and maintain the same sequence.

### Example Story 1 Highly recalled clauses for narrative of length 18 clauses

Original Story

1. My best friend pushed me into the pool.
2. This happened during my cousin’s wedding reception,
3. where everyone was dressed to impress.
4. And the reason it happened,
5. she spotted a bee hovering near my face,
6. -this was at a fancy hotel garden -
7. and she tried to save me from being stung.
8. And my aunt had just handed me a glass of champagne,
9. and I warned her to be careful.
10. ‘Course friends, y’know, they don’t always think things through.
11. So that’s when she gave me a little shove,
12. and I tumbled into the water.
13. When I resurfaced, gasping for air,
14. she just started laughing,
15. and she apologized profusely.
16. And… my beautiful dress was ruined,
17. and naturally, the first thing to do was to get out and dry off,
18. and my cousin just says, “Just about a few inches more,” she says, “and you’d have landed on the cake.”

Highly recalled clauses

*P*_*rec*_ = 0.857 1. My best friend pushed me into the pool.

*P*_*rec*_ = 0.893 2. This happened during my cousin’s wedding reception,

*P*_*rec*_ = 0.893 5. she spotted a bee hovering near my face,

*P*_*rec*_ = 0.661 16. And… my beautiful dress was ruined,

*P*_*rec*_ = 0.857 18. and my cousin just says, “Just about a few inches more,” she says, “and you’d have landed on the cake.”

### Example Story 2 Highly recalled clauses for narrative of length 54 clauses

Original Story

1. Let me share a story from when I was twenty-two years old, just fresh out of college
2. I was staying in a small town about five miles from here
3. I had just started my first job at a local bookstore
4. It was a quiet afternoon, and I was alone in the store
5. My grandpa, who had been bedridden for almost a year, was at home with my sister
6. She had to care for him daily, as he couldn’t move around on his own
7. There were only two of us siblings, my sister and me
8. My parents had gone out of town for the weekend
9. As I was shelving books, I suddenly heard a voice inside my head
10. It said, “If you go to the old church down the street and light a candle for your grandpa, he will get better
11. I looked around the bookstore, but no one was there
12. I continued with my work, trying to ignore the voice
13. It spoke to me again, repeating the same message
14. I looked around once more, but still found no one
15. I decided to take a break and walk to the church
16. I was only a short distance away, maybe a quarter-mile
17. As I approached the church, I hesitated for a moment
18. I was not a particularly religious person
19. I remembered my grandma always saying that prayers from a sincere heart could bring miracles
20. On the other hand, she also said that prayers without faith were useless
21. I admitted to myself that I was far from being a saint
22. I had my share of mistakes and misdeeds
23. I thought about my grandpa and how much he meant to our family
24. Finally, I went inside the church and found a candle
25. I knelt down in front of the altar
26. I said, “God, I don’t know how to pray properly
27. But I am asking you with all my heart to heal my grandpa
28. Please help him regain his strength and guide us in taking care of him
29. After saying my prayer, I returned to the bookstore
30. I resumed my work, shelving books
31. Suddenly, I felt a strong urge to go home
32. It was as if something was pulling me towards the house
33. I didn’t hear any voice this time
34. It was just an overwhelming feeling that I needed to go
35. I locked up the store and rushed home
36. To my surprise, I found my grandpa sitting in the living room, talking to my sister
37. He hadn’t been able to leave his bed for months without assistance
38. My sister told me that he had suddenly felt a surge of energy and was able to stand up and walk
39. That day marked the beginning of his recovery
40. He gradually regained his strength and independence
41. It was a turning point in our lives
42. We became more grateful for the time we had together
43. I often thought back to that day in the church
44. I wondered if my prayer had anything to do with my grandpa’s recovery
45. I couldn’t be sure, but it definitely changed my perspective on faith and the power of prayer
46. My grandpa lived for many more years after that incident
47. He celebrated his ninetieth birthday surrounded by family and friends
48. We cherished every moment we had with him
49. That experience happened almost twenty years ago
50. It remains a vivid memory in my mind
51. My grandpa passed away a few years later, but his spirit lives on in our hearts
52. Sometimes, when I’m feeling lost or uncertain, I think back to that day
53. I remember the power of faith and the miracles it can bring
54. And I am reminded that we are never truly alone, even when it seems like we are

Highly recalled clauses

*P*_*rec*_ = 0.710 1. Let me share a story from when I was twenty-two years old, just fresh out of college.

*P*_*rec*_ = 0.830 3. I had just started my first job at a local bookstore.

*P*_*rec*_ = 0.850 5. My grandpa, who had been bedridden for almost a year, was at home with my sister.

*P*_*rec*_ = 0.860 9. As I was shelving books, I suddenly heard a voice inside my head.

*P*_*rec*_ = 0.920 10. It said, “If you go to the old church down the street and light a candle for your grandpa, he will get better.”

*P*_*rec*_ = 0.720 15. I decided to take a break and walk to the church.

*P*_*rec*_ = 0.700 31. Suddenly, I felt a strong urge to go home.

*P*_*rec*_ = 0.900 36. To my surprise, I found my grandpa sitting in the living room, talking to my sister.

*P*_*rec*_ = 0.780 46. My grandpa lived for many more years after that incident.

## Appendix F. Narratives used in the study

The narratives that were generated by GPT-4 for this study used an existing segmentation into clauses. The original segmentation was taken from [Labov and Waletzky, 1966; Labov, 2013]. These segmentations were provided to the LLM to generate new variants, as detailed in appendix A.1. Below, we present all narratives that were presented to participants in our experiments. In the examples, the narratives are labeled by a prefix reflecting the source story, with the exception of “panic attack” which was created by GPT-3. The scrambled narratives are based on intact narrative (without “-scrambled” in the narrative name) where all clauses were shuffled. The “panic” story was generated based on the “boyscout” story from [Labov and Waletzky, 1966], Story 10 below. The story was generated using Prompt 6.

Story 3 and Story 4 below were generated using Prompt 1, with the Jacob Schissel story from [Labov, 2013] as a template.

### Example Story 4 Narrative schissel-v2-lake

1. My best friend pushed me into the lake.
2. This happened during our annual summer camping trip
3. and we were enjoying a peaceful afternoon.
4. The reason it all began,
5. she spotted a frog near the water’s edge or
6. -this was at our favorite spot in the woods -
7. and she dared me to catch it.
8. I had just sat down to eat my sandwich
9. and I told her I wasn’t interested.
10. ‘Course friends, y’know, they don’t always take no for an answer.
11. So that’s when she snuck up behind me
12. and gave me a little nudge.
13. When I emerged from the water, there was a frog on the shore,
14. she just scooped it up
15. and handed it to me.
16. And …; I couldn’t help but laugh – uncontrollably.
17. And naturally, the first thing to do was to get revenge,
18. and I said, “Just you wait,” I said, “you’ll get yours soon enough.”

Story 5 and Story 6 below were generated using Prompt 1, with the Charles Triplett story from [Labov, 2013] as a template.

### Example Story 5 Narrative triplett-v1-rookie

1. Back in the days when I was a rookie in the police force, the Chief was a veteran.
2. He had caught a notorious criminal –
3. or had been awarded for bravery.
4. But he had – a young daughter.
5. And in those days I was a fitness enthusiast.
6. And it seemed she was trying to impress me.
7. I never observed it.
8. Truth is I wasn’t very fond of her because she was –
9. She was a charming girl until she opened her mouth.
10. She was a chatterbox.
11. Goodness, she could talk.
12. Then she left a message one day saying she was going to run away because he was always scolding her about me.
13. He arrived at my apartment.
14. Big intimidating look on his face.
15. I managed to calm him down.
16. and proposed “Well we’ll search for her
17. and if we can’t locate her well you can – do whatever you think is right.”
18. I was strategizing.
19. So he accepted my offer.
20. And we went to where they found her scarf – near a park.
21. and we traced down a little more
22. and we couldn’t locate her.
23. And returned
24. – it was a police station –
25. she was sitting on a chair with a book in her hands.
26. She hadn’t run away.
27. But – nevertheless – that resolved the issue for that day.
28. But that evening the deputy, Frank Mitchell, said “You better transfer and leave because that old man never forgets anything once he gets it into his mind.”
29. And I did.
30. I transferred.
31. and left.
32. That was the end of my rookie year.

Story 7 and Story 8 below were generated using Prompt 1, with the Adolphus Hester story from [Labov, 2013] as a template.

### Example Story 6 Narrative triplett-v2-catlady

1. In the neighborhood I grew up in, Mrs. Baker was the resident cat lady.
2. She had adopted at least twenty –
3. or had given them homes.
4. But she had – a young grandson
5. and back then I was quite the tomboy.
6. And apparently, he was trying to befriend me.
7. I never caught on.
8. Truth is I didn’t take to him much because he had –
9. He was a charming lad until you saw his manners.
10. He had poor manners.
11. Goodness gracious he had poor manners.
12. Then he left a note one day saying he was going to run away because she was always nagging about me.
13. She came to my treehouse.
14. Big old tabby cat in tow.
15. I talked her down
16. and said “Well we’ll go find him
17. and if we can’t locate him, you can – go ahead and call the police if you need to.”
18. I was strategizing.
19. So she took my advice.
20. And we went to where they found his baseball cap – near the old mill
21. and we trailed a bit more
22. and we didn’t find a trace.
23. And returned
24. – it was a treehouse club –
25. he was sitting on a branch with a comic book in his hand.
26. He hadn’t run away.
27. But – however – that resolved it for that day.
28. But that evening the local sheriff, Officer Dawson said “You better lay low and stay out of sight because that woman never forgets anything once she gets it into her mind.”
29. And I did.
30. I laid low
31. and stayed out of sight.
32. That was the second time.

### Example Story 7 Narrative hester-v1-park

1. Let me share a story from when I was twenty-one years old
2. I was living in a small apartment near the city center
3. I had just started my first job
4. And uh I was working as a graphic designer
5. My father, who was recovering from a surgery, hadn’t been out in – in uh – well I’d say about eight months
6. He had to use a wheelchair to get around
7. The only person there with him during the day was a kind neighbor, a lady in her sixties
8. My younger sister was away at college –
9. so it was just my dad and the neighbor lady at home during the day
10. One day, I got this strong feeling while I was at work
11. It told me: “If you go to the nearby park during your lunch break and pray for your father’s recovery, he’ll start to improve
12. I looked around my office
13. I didn’t see anyone nearby
14. So, I went to the park during my break
15. and found a quiet spot there
16. That feeling came back again
17. I looked around once more
18. I still didn’t see anyone nearby
19. but I decided to take a chance
20. I wasn’t too far from my office, maybe a ten-minute walk
21. I found a bench to sit on and closed my eyes – uh – uh –
22. that feeling came back once more
23. I just took a deep breath
24. I was sitting under a big oak tree now
25. I looked up at the branches above me
26. Nobody was around
27. And uh I said to myself, well now,
28. I had been taught that prayers can make a difference
29. And that it doesn’t hurt to try
30. I knew I wasn’t perfect, but I wanted to help my dad
31. I’d admit that to anyone
32. But I was willing to give it a try
33. I looked around one more time
34. and then I closed my eyes
35. and said a quiet prayer
36. I whispered, “God, I don’t know if you’re listening
37. But I ask you to please help my father recover
38. So he can regain his independence and enjoy life again
39. I opened my eyes
40. and went back to work
41. Suddenly, I received a text message from my father
42. It said, “I just took a few steps with my walker
43. I – I couldn’t believe it
44. It felt like a sign that my prayer had been heard
45. I rushed home after work
46. and saw my father standing in the living room with the help of his walker
47. He hadn’t been able to do that in months without assistance
48. And there he was, taking small steps
49. That was the turning point in his recovery
50. He continued to improve
51. and regained most of his mobility within a year
52. It’s been, what, six or seven years since then?
53. My father is now able to walk without any assistance
54. He’s living life to the fullest, even in his seventies

### Example Story 8 Narrative hester-v2-church

1. Let me share a story from when I was twenty-two years old, just fresh out of college.
2. I was staying in a small town about five miles from here.
3. I had just started my first job at a local bookstore.
4. It was a quiet afternoon, and I was alone in the store.
5. My grandpa, who had been bedridden for almost a year, was at home with my sister.
6. She had to care for him daily, as he couldn’t move around on his own.
7. There were only two of us siblings, my sister and me.
8. My parents had gone out of town for the weekend.
9. As I was shelving books, I suddenly heard a voice inside my head.
10. It said, “If you go to the old church down the street and light a candle for your grandpa, he will get better.”
11. I looked around the bookstore, but no one was there.
12. I continued with my work, trying to ignore the voice.
13. It spoke to me again, repeating the same message.
14. I looked around once more, but still found no one.
15. I decided to take a break and walk to the church.
16. I was only a short distance away, maybe a quarter-mile.
17. As I approached the church, I hesitated for a moment.
18. I was not a particularly religious person.
19. I remembered my grandma always saying that prayers from a sincere heart could bring miracles.
20. On the other hand, she also said that prayers without faith were useless.
21. I admitted to myself that I was far from being a saint.
22. I had my share of mistakes and misdeeds.
23. I thought about my grandpa and how much he meant to our family.
24. Finally, I went inside the church and found a candle.
25. I knelt down in front of the altar.
26. I said, “God, I don’t know how to pray properly.
27. But I am asking you with all my heart to heal my grandpa.
28. Please help him regain his strength and guide us in taking care of him.”
29. After saying my prayer, I returned to the bookstore.
30. I resumed my work, shelving books.
31. Suddenly, I felt a strong urge to go home.
32. It was as if something was pulling me towards the house.
33. I didn’t hear any voice this time.
34. It was just an overwhelming feeling that I needed to go.
35. I locked up the store and rushed home.
36. To my surprise, I found my grandpa sitting in the living room, talking to my sister.
37. He hadn’t been able to leave his bed for months without assistance.
38. My sister told me that he had suddenly felt a surge of energy and was able to stand up and walk.
39. That day marked the beginning of his recovery.
40. He gradually regained his strength and independence.
41. It was a turning point in our lives.
42. We became more grateful for the time we had together.
43. I often thought back to that day in the church.
44. I wondered if my prayer had anything to do with my grandpa’s recovery.
45. I couldn’t be sure, but it definitely changed my perspective on faith and the power of prayer.
46. My grandpa lived for many more years after that incident.
47. He celebrated his ninetieth birthday surrounded by family and friends.
48. We cherished every moment we had with him.
49. That experience happened almost twenty years ago.
50. It remains a vivid memory in my mind.
51. My grandpa passed away a few years later, but his spirit lives on in our hearts.
52. Sometimes, when I’m feeling lost or uncertain, I think back to that day.
53. I remember the power of faith and the miracles it can bring.
54. And I am reminded that we are never truly alone, even when it seems like we are.

Story 9 below is taken from the story of Gloria Stein (Ch. 7) in [Labov, 2013].

### Example Story 9 Narrative stein

1. We moved down to Third and Wharton
2. We had gotten moved
3. but it was like – you remember I told you my father drank,
4. but it was still day,
5. and after he had gotten in and put the furniture where it was supposed to go, my mother left, to go to market to get some food to bring into the house
6. And my father went to take the horses – take the team of horses back
7. But he didn’t come home
8. He – stopped somewhere on – I dunno,
9. but he – how he – why he wasn’t in the house
10. And of course I was left there with the kids
11. Well no sooner did my mother get out of the house – my father went first – then – oh it was such an awful crash came through the window
12. And I – and I looked and it was a big stone
13. And then of course they – they – well they stoned the house
14. And I had sense enough to get the kids into the stairway so that they wouldn’t get hurt by flyin’ glass
15. And I said to them, “Now you stay there
16. Don’t move ‘cause nothing will reach you here
17. And I’m gonna go and see if I can find mom” –
18. (We call our mother mom
19. And they were crying
20. and carryin’ on
21. I was scared to death,
22. I really was frightened out of my wits
23. And I didn’t know what to do,
24. but I remember saying a prayer as I went out the front door because I didn’t know what was gonna hit me when I went out that door
25. And I ran –
26. I was ten years old at the time this happened –
27. I ran to Fourth street, because I knew my mother wa-
28. They had push carts in those days, on Fourth street
29. And it was a Jewish neighborhood
30. and they – they stayed open until – until all hours
31. Because they would, you know, however much money they could get they wanted
32. So uh that’s where she was going to do her shopping
33. And I finally found my mother
34. I did find her
35. And while I was trying to tell her what had happened
36. a policeman went by
37. And my voice was such that you could hardly make out what I was saying
38. I was so nervous and – and – my voice was shaking so,
39. and he wanted to find out what ha-what was wrong with me
40. And then when he understood what the situation was he turned to my mother
41. and says, “This is a bad neighborhood
42. It’s Irish
43. And he said, “They’re known to do things like that,
44. Better move out of there
45. My mother said, “I just moved in, how am I gonna move out?”
46. He says, “I dunno
47. He says, “But they don’t want any black people in there
48. An’ he didn’t say black ‘cause they didn’t use black then
49. But “they don’t want anybody in there
50. So uh, she said, “Well you’re the law
51. You’re supposed to protect us
52. He says, “I can’t do anything with it
53. He was Irish himself
54. “I can’t do anything about it
55. So my mother said um, “Where is the uh precinct you work from?”
56. And he told her
57. I forgot now
58. I was right there with her
59. but I’ve forgotten where he told her
60. So uh she said, “Well if you can’t do anything to help me, I’ll go see who – if I can’t find someone else to help me
61. Well in those days black people were so cowed by circumstances an’ uniforms and white people, you know, they uh many times didn’t assert themselves
62. But thankfully my mother was not one of those kind
63. And she – well I always thought of her as an educated person which she was because she was the only one I knew who had completed high school
64. She went to Lackawanna high school
65. And this was – let me see what year was this happening – well it must have been nineteen twenty-five, so you can see what – what the times were
66. And we went – she took me, and we went to the police station
67. And they also had police wagons with horses attached to them
68. So my mother said, to the captain, “What are you going to do about it?”
69. And he says, “Well lady there’s nothing we can do about it
70. And she says, “You mean to tell me that I have to live under those circumstances and there’s nothing that you can do about it?”
71. She says – he says, “Well uh the only thing I can tell you to do is to move
72. He says, “Because” – he said to me, “Did you see anyone?”
73. And I said, “No
74. He says, “Well how can – how can you” – to my mother – “how can you do anything if you don’t know who – who to put the blame on?”
75. An’ he says, “I’m sure there was more than one person
76. So my mother said, “Well I don’t know what I’m going to do
77. She says, “But one thing I know I’m not going to do is move
78. They took all we had to, to pay the rent
79. In that house you had to pay in advance
80. And she said, “If they didn’t want me there they should not have rented the house to me
81. Now the real estate agent should be accountable
82. So anyhow, she said to him, “Well I know one thing you can do
83. And he says, “What’s that?”
84. She says, “You can take me home
85. He said, “Take you home?”
86. She said, “Yes in the police patrol
87. So – I guess they figured “Let’s get this lady out of here because she’s talking loud and long
88. And they had never come up with anybody like that
89. So they put us in the police patrol
90. and they took us home
91. We weren’t far from – from where it was – maybe Fifth street – from where we had moved
92. And – course with the horses and the police patrol, and the clang –
93. they had a bell that clanged
94. you know they told you about the trolley that went “Clang
95. Well this police patrol that bell they had, it went “Clang clang clang
96. And um all the neighbors came running out to see what was happening
97. And of course they saw the man came around the back –
98. I can see it right now – from where he was sitting up front there where the horses came around,
99. and opened the door in the back
100. And they had a long door in the back where you had to come down steps to get out the back of the – of this police patrol
101. Well we got out of that police patrol –
102. Before the patrol ever left my mother turned
103. and faced all those people that were standing around,
104. and she says, “Why you’re doing this to me I don’t know
105. She says, “I know that I am different from you, color wise,
106. but in every other respect I’m just the same as you are,
107. and all I’m trying to do is provide a home for my family
108. She told them that they had rented the house,
109. she told them how much money she had paid,
110. she had told them that was all the money she had
111. And they had paid that rent for a month
112. She knew she was going to be there for the month
113. She did not know beyond the month
114. Whether they’d have enough money to pay it for the next month
115. But she knew they were going to be there for the month
116. She said, “It is winter time and it is cold
117. And I cannot afford to be buying windows
118. “Now” she said, “Whoever threw the stones I don’t know
119. But you can all see what has happened
120. And now I’m telling you that we’re going to be here for a month
121. So you might just as well make up your mind to that
122. If you don’t want me here then you people get together and give me enough money to move somewhere else, because I do not have it
123. Well of course you know they didn’t do that
124. They probably didn’t have any money either
125. So in the meantime – not in the meantime but during, during the period of time that we were there, my father became sick
126. And the neighbors were as nice as anybody could ever have been
127. When there was trouble they responded just like anyone else
128. But that was our welcoming to the neighborhood
129. And it was terrible
130. and it certainly was frightening for me

Below, Story 10 is taken directly from [Labov and Waletzky, 1966]. Story 11 is the scrambled version used in experiments.

### Example Story 10 Narrative boyscout

1. Yeah, I was in the boy scouts at the time.
2. And we was doing the 50-yard dash,
3. racing,
4. but we was at the pier, marked off,
5. and so we was doing the 50-yard dash.
6. There was about 8 or 9 of us, you know, going down, coming back.
7. And, going down the third time, I caught cramps
8. and I started yelling “Help!”,
9. but the fellows didn’t believe me, you know.
10. They thought I was just trying to catch up, because I was going on or slowing down.
11. So all of them kept going.
12. They leave me.
13. And so I started going down.
14. Scoutmaster was up there.
15. He was watching me.
16. But he didn’t pay me no attention either.
17. And for no reason at all there was another guy, who had just walked up that minute…
18. He just jumped over
19. and grabbed me.

### Example Story 11 Narrative boyscout-scrambled

1. They leave me.
2. racing,
3. Yeah, I was in the boy scouts at the time.
4. but we was at the pier, marked off,
5. and so we was doing the 50-yard dash.
6. And we was doing the 50-yard dash,
7. but the fellows didn’t believe me, you know.
8. and I started yelling “Help!”,
9. and grabbed me.
10. Scoutmaster was up there.
11. So all of them kept going.
12. And so I started going down.
13. There was about 8 or 9 of us, you know, going down, coming back.
14. He just jumped over
15. They thought I was just trying to catch up, because I was going on or slowing down.
16. And, going down the third time, I caught cramps
17. And for no reason at all there was another guy, who had just walked up that minute…
18. He was watching me.
19. But he didn’t pay me no attention either

Story 12, Story 13, and Story 14 are the scrambled versions of Story 5,Story 8, and Story 9, respectively. The scrambled is done by taking a random permutation of the clauses. The numbering in the scrambled versions reflects the presentation order.

### Example Story 12 Narrative triplett-v1-rookie-scrambled

1. and we traced down a little more
2. and left.
3. And I did.
4. I never observed it.
5. I was strategizing.
6. Big intimidating look on his face.
7. and proposed “Well we’ll search for her
8. Back in the days when I was a rookie in the police force, the Chief was a veteran.
9. and we couldn’t locate her.
10. And in those days I was a fitness enthusiast.
11. And we went to where they found her scarf – near a park.
12. But – nevertheless – that resolved the issue for that day.
13. That was the end of my rookie year.
14. And it seemed she was trying to impress me.
15. or had been awarded for bravery.
16. He had caught a notorious criminal –
17. She hadn’t run away.
18. She was a chatterbox.
19. I managed to calm him down.
20. – it was a police station –
21. And returned
22. Truth is I wasn’t very fond of her because she was –
23. Then she left a message one day saying she was going to run away because he was always scolding her about me.
24. But that evening the deputy, Frank Mitchell, said “You better transfer and leave because that old man never forgets anything once he gets it into his mind.”
25. I transferred.
26. But he had – a young daughter.
27. Goodness, she could talk.
28. So he accepted my offer.
29. She was a charming girl until she opened her mouth.
30. she was sitting on a chair with a book in her hands.
31. He arrived at my apartment.
32. and if we can’t locate her well you can – do whatever you think is right.”

### Example Story 13 Narrative hester-v2-church-scrambled

1. I remembered my grandma always saying that prayers from a sincere heart could bring miracles
2. Finally, I went inside the church and found a candle
3. As I was shelving books, I suddenly heard a voice inside my head
4. I remember the power of faith and the miracles it can bring
5. It was a turning point in our lives
6. That experience happened almost twenty years ago
7. I was only a short distance away, maybe a quarter-mile
8. My grandpa lived for many more years after that incident
9. I said, “God, I don’t know how to pray properly
10. And I am reminded that we are never truly alone, even when it seems like we are
11. It spoke to me again, repeating the same message
12. I was not a particularly religious person
13. Suddenly, I felt a strong urge to go home
14. I looked around the bookstore, but no one was there
15. I was staying in a small town about five miles from here
16. My sister told me that he had suddenly felt a surge of energy and was able to stand up and walk
17. Sometimes, when I’m feeling lost or uncertain, I think back to that day
18. I wondered if my prayer had anything to do with my grandpa’s recovery
19. Let me share a story from when I was twenty-two years old, just fresh out of college
20. My parents had gone out of town for the weekend
21. I locked up the store and rushed home
22. It was as if something was pulling me towards the house
23. I admitted to myself that I was far from being a saint
24. I had just started my first job at a local bookstore
25. As I approached the church, I hesitated for a moment
26. She had to care for him daily, as he couldn’t move around on his own
27. It was a quiet afternoon, and I was alone in the store
28. We cherished every moment we had with him
29. My grandpa, who had been bedridden for almost a year, was at home with my sister
30. I had my share of mistakes and misdeeds
31. He hadn’t been able to leave his bed for months without assistance
32. There were only two of us siblings, my sister and me
33. I didn’t hear any voice this time
34. But I am asking you with all my heart to heal my grandpa
35. My grandpa passed away a few years later, but his spirit lives on in our hearts
36. I looked around once more, but still found no one
37. He celebrated his ninetieth birthday surrounded by family and friends
38. That day marked the beginning of his recovery
39. We became more grateful for the time we had together
40. I continued with my work, trying to ignore the voice
41. It was just an overwhelming feeling that I needed to go
42. I knelt down in front of the altar
43. I thought about my grandpa and how much he meant to our family
44. Please help him regain his strength and guide us in taking care of him
45. To my surprise, I found my grandpa sitting in the living room, talking to my sister
46. I often thought back to that day in the church
47. I decided to take a break and walk to the church
48. I couldn’t be sure, but it definitely changed my perspective on faith and the power of prayer
49. After saying my prayer, I returned to the bookstore

### Example Story 14 Narrative stein-scrambled

1. She says, “But one thing I know I’m not going to do is move. Because I have no money to move with.”
2. And they also had police wagons with horses attached to them. Believe it or not in those days.
3. they had a bell that clanged. Clang clang clang –
4. And I didn’t know what to do,
5. But thankfully my mother was not one of those kind.
6. He says, “I dunno.”
7. We weren’t far from – from where it was – maybe Fifth street – from where we had moved.
8. In that house you had to pay in advance.
9. She says – he says, “Well uh the only thing I can tell you to do is to move.”
10. Don’t move ‘cause nothing will reach you here.
11. but in every other respect I’m just the same as you are,
12. So uh, she said, “Well you’re the law.
13. And they had never come up with anybody like that.
14. and opened the door in the back.
15. And I cannot afford to be buying windows.”
16. She says, “I know that I am different from you, color wise,
17. I was so nervous and – and – my voice was shaking so,
18. An’ he didn’t say black ‘cause they didn’t use black then.
19. I can see it right now – from where he was sitting up front there where the horses came around,
20. (We call our mother mom.)
21. I was right there with her
22. So my mother said um, “Where is the uh precinct you work from?”
23. And she says, “You mean to tell me that I have to live under those circumstances and there’s nothing that you can do about it?”
24. She says, “You can take me home.”
25. And I said, “No.”
26. And it was terrible
27. But that was our welcoming to the neighborhood.
28. And now I’m telling you that we’re going to be here for a month.
29. And she – well I always thought of her as an educated person which she was because she was the only one I knew who had completed high school.
30. He says, “Because” – he said to me, “Did you see anyone?”
31. They had push carts in those days, on Fourth street.
32. She said, “It is winter time and it is cold.
33. Before the patrol ever left my mother turned
34. So you might just as well make up your mind to that.
35. but I’ve forgotten where he told her.
36. And he told her.
37. And I finally found my mother.
38. Well no sooner did my mother get out of the house – my father went first – then – oh it was such an awful crash came through the window.
39. So uh that’s where she was going to do her shopping. 40. But “they don’t want anybody in there.”
40. I was scared to death,
41. And I had sense enough to get the kids into the stairway so that they wouldn’t get hurt by flyin’ glass.
42. And he said, “They’re known to do things like that,
43. An’ he says, “I’m sure there was more than one person.”
44. It’s Irish.”
45. but I remember saying a prayer as I went out the front door because I didn’t know what was gonna hit me when I went out that door.
46. Whether they’d have enough money to pay it for the next month.
47. And this was – let me see what year was this happening – well it must have been nineteen twenty-five, so you can see what – what the times were.
48. And I’m gonna go and see if I can find mom” –
49. Better move out of there.”
50. I forgot now.
51. Because they would, you know, however much money they could get they wanted.
52. and she says, “Why you’re doing this to me I don’t know.”
53. So anyhow, she said to him, “Well I know one thing you can do.”
54. and carryin’ on.
55. So – I guess they figured “Let’s get this lady out of here because she’s talking loud and long.”
56. And I – and I looked and it was a big stone.
57. you know they told you about the trolley that went “Clang.”
58. And they had paid that rent for a month.
59. He said, “Take you home?”
60. He says, “Well how can – how can you” – to my mother – “how can you do anything if you don’t know who – who to put the blame on?”
61. I did find her.
62. I really was frightened out of my wits.
63. but he – how he – why he wasn’t in the house.
64. She knew she was going to be there for the month.
65. a policeman went by.
66. Now the real estate agent should be accountable.”
67. And while I was trying to tell her what had happened
68. And we went – she took me, and we went to the police station.
69. Well of course you know they didn’t do that.
70. And of course they saw the man came around the back –
71. And my voice was such that you could hardly make out what I was saying.
72. and they – they stayed open until – until all hours.
73. So in the meantime – not in the meantime but during, during the period of time that we were there, my father became sick.
74. And he says, “What’s that?”
75. He was Irish himself.
76. We moved down to Third and Wharton.
77. He says, “I can’t do anything with it.”
78. So my mother said, “Well I don’t know what I’m going to do.”
79. she told them how much money she had paid,
80. “Now” she said, “Whoever threw the stones I don’t know.
81. You’re supposed to protect us.”
82. So my mother said, to the captain, “What are you going to do about it?”
83. and faced all those people that were standing around,
84. And the neighbors were as nice as anybody could ever have been.
85. They probably didn’t have any money either.
86. Well this police patrol that bell they had, it went “Clang clang clang.”
87. He says, “But they don’t want any black people in there.”
88. But you can all see what has happened.
89. And she said, “If they didn’t want me there they should not have rented the house to me.
90. And I ran –
91. and it certainly was frightening for me.
92. And he says, “Well lady there’s nothing we can do about it.”
93. and they took us home.
94. I ran to Fourth street, because I knew my mother wa-
95. Well we got out of that police patrol –
96. “I can’t do anything about it.”
97. He – stopped somewhere on – I dunno,
98. She said, “Yes in the police patrol.”
99. And they had a long door in the back where you had to come down steps to get out the back of the – of this police patrol.
100. And then of course they – they – well they stoned the house.
101. So they put us in the police patrol
102. They took all we had to, to pay the rent.
103. We had gotten moved
104. and all I’m trying to do is provide a home for my family.”
105. But she knew they were going to be there for the month.
106. and after he had gotten in and put the furniture where it was supposed to go, my mother left, to go to market to get some food to bring into the house.
107. And they were crying
108. And um all the neighbors came running out to see what was happening.
109. When there was trouble they responded just like anyone else.
110. She told them that they had rented the house,
111. If you don’t want me here then you people get together and give me enough money to move somewhere else, because I do not have it.”
112. And – course with the horses and the police patrol, and the clang –
113. And I said to them, “Now you stay there.
114. And then when he understood what the situation was he turned to my mother
115. she had told them that was all the money she had.
116. She went to Lackawanna high school.
117. I was ten years old at the time this happened –
118. And of course I was left there with the kids.
119. and he wanted to find out what ha-what was wrong with me.
120. And my father went to take the horses – take the team of horses back.
121. and says, “This is a bad neighborhood.
122. but it was still day,
123. And it was a Jewish neighborhood
124. but it was like – you remember I told you my father drank,
125. But he didn’t come home.
126. So uh she said, “Well if you can’t do anything to help me, I’ll go see who – if I can’t find someone else to help me.”
127. She did not know beyond the month.
128. Well in those days black people were so cowed by circumstances an’ uniforms and white people, you know, they uh many times didn’t assert themselves.
129. My mother said, “I just moved in, how am I gonna move out?”

## Appendix G. List of Nouns

The following list of 32 nouns in exactly this order was presented to each participant in the Random List of Nouns experiment.

### Example 1

‘FAN’, ‘POLICE’, ‘SURVEY’, ‘COFFEE’, ‘ISLAND’, ‘CALF’, ‘ACID’, ‘SUITCASE’, ‘TIDE’, ‘BULLET’, ‘BUILDING’, ‘TELEVISION’, ‘QUEEN’, ‘BEDROOM’, ‘LIGHTNING’, ‘MOUNTAIN’, ‘LEMON’, ‘SEAT’, ‘CITY’, ‘VICTIM’, ‘LIST’, ‘CHAIN’, ‘STEM’, ‘ATOM’, ‘STAKE’, ‘DEER’, ‘OPERATOR’, ‘CROSS’, ‘FLOOR’, ‘INFANT’, ‘INDIAN’, ‘PASSAGE’

Narrative Memory A PREPRINT

## Appendix H. Story used for LLM reliability assessment

The following narrative was used for comparison of recall performance evaluation by humans versus GPT in fig. 1.

The story was generated with an older version of GPT (`text-davinci-003`) using the following prompt seeded by the boyscouts narrative (Story 10, in paragraph form without the numbering):

### Prompt 5

**Panic Attack Story Generation**

~~~
This is an oral narrative about a life-threatening event:
{ story }
Create a different, longer and more detailed oral narrative with a similar narration style about another life-threatening event.
~~~

yielding the following narrative:

### Argument 5

**Panic Attack Narrative**

Well, I was at work one day and I was doing my usual job - working on the warehouse floor, stocking boxes and organizing merchandise. I had been working for a few hours and it was getting late. I had been feeling a bit strange, a bit lightheaded, but chalked it up to hunger and not having eaten yet that day. But then, as I was walking around the warehouse returning items to their proper places, I started to feel a strange tightness and pressure in my chest. I stopped what I was doing and leaned against a nearby wall for a bit, but the pressure only got stronger. And I could feel my heart pounding in my chest. Next thing I know, I started seeing spots and I could feel sweat dripping down my forehead and my arms. I tried to take some deep breaths, but I just couldn’t catch my breath. I was really scared and didn’t know what to do. Suddenly I heard a voice coming from across the warehouse. It was the manager, who had seen me and called out to me - it seems someone had noticed me and was checking to see if I was alright. I tried to yell back that something was wrong, but I couldn’t find my voice. The manager started walking towards me and I could feel my world going dark. I started to drop to the ground, but luckily the manager got to me in time and caught me just before my legs gave out. He took me to the staff room and sat me down in a chair. He asked me questions about what was wrong and what happened. He got me a glass of cold water and told me to rest for a bit. After a few minutes, the pressure in my chest slowly eased up and I started to feel better. It turned out I had been having a panic attack, but I escaped from the situation with the help of the quick action of the manager. It was never a fun experience, but it really could have been much worse, and it taught me to check in with myself and make sure to properly rest and eat to avoid further episodes.

The story was subsequently segmented using the prompt:

### Prompt 6

**Segmentation Prompt**

~~~
This is the original story:
{Argument 5}
Split the sentences of the original story into the maximum number of small pieces, such that each piece contains minimal information.
~~~

The segmentation that comes up is the following:

### Completion 5

**Panic Attack Segmentation**

1. I was at work.
2. I was working on the warehouse floor.
3. I was stocking boxes and organizing merchandise.
4. I had been working for a few hours.
5. It was getting late.
6. I had been feeling a bit strange.
7. I was feeling lightheaded.
8. I chalked it up to hunger and not having eaten yet that day.
9. I was walking around the warehouse returning items to their proper places.
10. I started to feel a strange tightness and pressure in my chest.
11. I stopped what I was doing.
12. I leaned against a nearby wall.
13. The pressure only got stronger.
14. I could feel my heart pounding in my chest.
15. I started seeing spots.
16. I could feel sweat dripping down my forehead and my arms.
17. I tried to take some deep breaths.
18. I couldn’t catch my breath.
19. I was really scared.
20. I didn’t know what to do.
21. I heard a voice coming from across the warehouse.
22. It was the manager.
23. Someone had noticed me.
24. The manager was checking to see if I was alright.
25. I tried to yell back that something was wrong.
26. I couldn’t find my voice.
27. The manager started walking towards me.
28. I could feel my world going dark.
29. I started to drop to the ground.
30. The manager got to me in time.
31. He caught me just before my legs gave out.
32. He took me to the staff room.
33. He sat me down in a chair.
34. He asked me questions about what was wrong and what happened.
35. He got me a glass of cold water.
36. He told me to rest for a bit.
37. The pressure in my chest slowly eased up.
38. I started to feel better.
39. It turned out I had been having a panic attack.
40. I escaped from the situation with the help of the quick action of the manager.
41. It was never a fun experience.
42. It really could have been much worse.
43. It taught me to check in with myself.
44. It taught me to properly rest and eat to avoid further episodes.

Note that the segmentation in this particular case is not verbatim, the clauses have been transformed to first person while retaining most important information. If a verbatim segmentation is desired however, it can be achieved by utilizing the same prompt with `gpt-4` although the number of clauses will be slightly different.

## Footnotes

2 The narratives ‘boyscout’ and ‘stein’ are taken directly from these references. The rest are generated using templates.

